# Stochastic coding: a conserved feature of odor representations and its implications for odor discrimination

**DOI:** 10.1101/2023.06.27.546757

**Authors:** Shyam Srinivasan, Simon Daste, Mehrab Modi, Glenn Turner, Alexander Fleischmann, Saket Navlakha

## Abstract

Sparse coding is thought to improve discrimination of sensory stimuli by reducing overlap between their representations. Two factors, however, can offset sparse coding’s advantages. Similar sensory stimuli have significant overlap, and responses vary across trials. To elucidate the effect of these two factors, we analyzed odor responses in the fly and mouse olfactory regions implicated in learning and discrimination — the Mushroom Body (MB) and the Piriform Cortex (PCx). In both species, we show that neuronal responses fall along a continuum from extremely reliable across trials to extremely variable or stochastic. Computationally, we show that the range of observed variability arises from probabilistic synapses in inhibitory feedback connections within central circuits rather than sensory noise, as is traditionally assumed. We propose this coding scheme to be advantageous for coarse– and fine-odor discrimination. More reliable cells enable quick discrimination between dissimilar odors. For similar odors, however, these cells overlap, and do not provide distinguishing information. By contrast, more unreliable cells are decorrelated for similar odors, providing distinguishing information, though this requires extended training with more trials. Overall, we have uncovered a stochastic coding scheme that is conserved in vertebrates and invertebrates, and we identify a candidate mechanism, based on variability in a winner-take-all inhibitory circuit, that improves discrimination with training.

## Introduction

Sparse coding has emerged as an important principle of neural computation (Field, 1994; Barth & Poulet, 2012). Sparse coding can improve discrimination by reducing overlap between representations (Lin et al., 2014) and leads to more efficient computation because fewer active neurons implies less energy expenditure (Attwell & Laughlin, 2001). Sparse coding has been observed across sensory modalities, ranging from vision and audition (Vinje & Gallant, 2000) to olfaction, and across species, from vertebrates (Stettler & Axel, 2009; Poo & Isaacson, 2009) to invertebrates (Turner et al., 2008; Campbell et al., 2013; Lin et al., 2014; Perez-Orive et al., 2002; Endo et al., 2020).

A closer examination of stimulus responses, however, poses a challenge for our understanding of sparse coding’s role in the brain. Most studies report averaged or thresholded stimulus responses across trials, which result in an underestimate of the responsive population. For example, in the olfactory system of fruit flies, the percentage of higher order neurons that respond to an odor can be up to 15% per trial (Honegger et al., 2011; Campbell et al., 2013), as opposed to 5% if averaged and thresholded across trials. Similarly, in the mammalian olfactory system, at least 16% of the coding population responds per trial in rats (Miura et al., 2012) and about 20% responds in mice (Roland et al., 2017), up from 10% (Stettler & Axel, 2009) when averaged across trials. The percentage response in anesthetized mice is less, although only slightly, than in awake ones (Bolding et al., 2020). The discrepancy in perceived sparsity as calculated from multiple trials versus single trials is relevant as one can envision real world scenarios, e.g., sensing a predator’s presence, that might not afford the luxury of multiple trials for decision-making. Therefore, explanations of animal behavior would be incomplete without taking into account the lower than perceived (Spanne & Jorntell, 2015) sparsity of sensory responses.

Neuroscientists must contend with a second issue in trying to explain cognitive processes downstream of peripheral sensory coding: variability in neural responses across trials. Trial-to-trial variability is ubiquitous in the brain. Amongst sensory regions, it is observed in the visual system (Carandini, 2004; Arazi et al., 2017; Sadagopan & Ferster, 2012), the auditory system (Lins-Ribeiro et al., 2020), the whisker thalamus (Scaglione et al., 2011), and the motor system (Mandairon & Linster, 2009; Wu et al., 2014). Within learning and memory systems, it is observed in the hippocampus and the entorhinal cortex (Prerau & Eden, 2018), the prefrontal cortex (Hussar & Pasternak, 2010), and the basal ganglia (Tumer & Brainard, 2007; Woolley & Kao, 2015). Such widespread observations of variability begs the question, if animals must make decisions based on a single exposure of a sensory stimulus, how can they produce robust decisions when stimulus representations might differ across exposures (or trials)?

To understand how sparsity level and variability affect decision making, three questions need to be resolved. First, how many cells respond consistently (i.e., across most trials) and how many total cells respond? Second, are there differences in the information encoded by these two populations? Third, what are the mechanisms that produce trial-to-trial variability? Traditionally, variability is viewed as a byproduct of sensory-level noise and is thus a “nuisance” the brain must contend with. Analyses that threshold or average responses across trials effectively support this view. In contrast, it has been noted that variability can be beneficial in some situations (Faisal et al., 2008; Ermentrout et al., 2008), suggesting that variability may be intrinsic to the circuit.

Our goal in this study is to characterize sparsity and variability of odor responses in the fly mushroom body (MB) and the mouse piriform cortex (PCx), and to understand the consequences of these factors towards discrimination and learning. The olfactory system is attractive to investigate these questions because it exhibits both sparsity and variability. In addition, as MB and PCx are only 2 synapses from the external environment, circuit anatomy and physiology linking sensory responses to neural coding and behavior are well defined, particularly in flies (Aso et al., 2014a; Zheng et al., 2018; Modi et al., 2020) and conserved in mammals (Bekkers & Suzuki, 2013; Neville & Haberly, 2004; Giessel & Datta, 2014). We show, in both flies (using existing MB data (Campbell et al., 2013)) and mice (using new PCx data collected for this study), that cells responding to an odor fall into two classes: a small set of cells have large responses and are reliably activated across trials, and a larger set of cells have smaller responses and are unreliably activated across trials. We show that reliable cell patterns can better decode odor identity than unreliable cells, suggesting that reliable cells represent a more stable (i.e., less stochastic) component of the odor code. However, we show that reliable cells by themselves have high overlap for similar odors, whereas unreliable cells do not. Thus, while reliable cells can easily distinguish dissimilar odors, we propose that unreliable cells can, with training, be used to distinguish very similar odors. Finally, using a computational model of the olfactory circuit, we show that the variability observed experimentally far exceeds what would be expected under reasonable levels of sensory noise, and that synaptic variability in the winner-take-all (WTA) inhibitory circuit is a more likely mechanism for generating stochastic codes.

## Results

### Basic anatomy and physiology of the olfactory circuit

In flies and mice, the olfactory circuit comprises three stages (Fig. 1A), with key elements of the circuit architecture being conserved across species. In stage 1, odor information is captured by olfactory sensory neurons (OSNs) in the mammalian nose (Mombaerts et al., 1996; Buck & Axel, 1991; Zhang & Firestein, 2009) or fly antenna and maxillary palp (Vosshall et al., 2000; Wilson, 2013), where each OSN type expresses a single receptor type that preferentially responds to a particular chemical class. There are about 1000 OSN types in mice and 50 types in flies, which are responsible for sensing the vast space of possible chemical compounds. Thus, odors are encoded as a combinatorial code; i.e., every odor is represented by a unique combination of neurons that respond to that odor (Stevens, 2015, 2016; Malnic et al., 1999). In stage 2, odor information captured by OSNs is transferred to glomeruli in the olfactory bulb of mammals (Murthy, 2011; Vassar et al., 1994; Ressler et al., 1994) or antennal lobe of flies (Wilson, 2013). Glomerular encoding is modified by a lateral inhibition circuit, which increases the reliability of the odor representation and normalizes its output range (Bhandawat et al., 2007; Carandini & Heeger, 2012). In stage 3, glomeruli, through projection neurons, pass odor information to piriform cortex (PCx) cells in mice or mushroom body (MB) Kenyon cells (KCs) in flies (Bekkers & Suzuki, 2013; Modi et al., 2020). In flies, a winner-take-all (WTA) circuit sparsens KC odor responses (Fig. 1A); all KCs activate a giant inhibitory neuron (called the Anterior Paired Lateral neuron), which then negatively feeds back to suppress the activity of less responsive KCs (Lin et al., 2014). PCx also contains a WTA circuit, wherein PCx principal cells activate inhibitory neurons in layers 2 and 3 of PCx, which then negatively feedback to suppress less responsive PCx cells (Bekkers & Suzuki, 2013; Franks et al., 2011; Bolding & Franks, 2018). The sparse encoding of the odor in the third stage (MB or PCx) is used downstream by the animal for olfactory discrimination and learning (Heisenberg, 2003; Choi et al., 2011; Modi et al., 2020).

**Figure 1:**
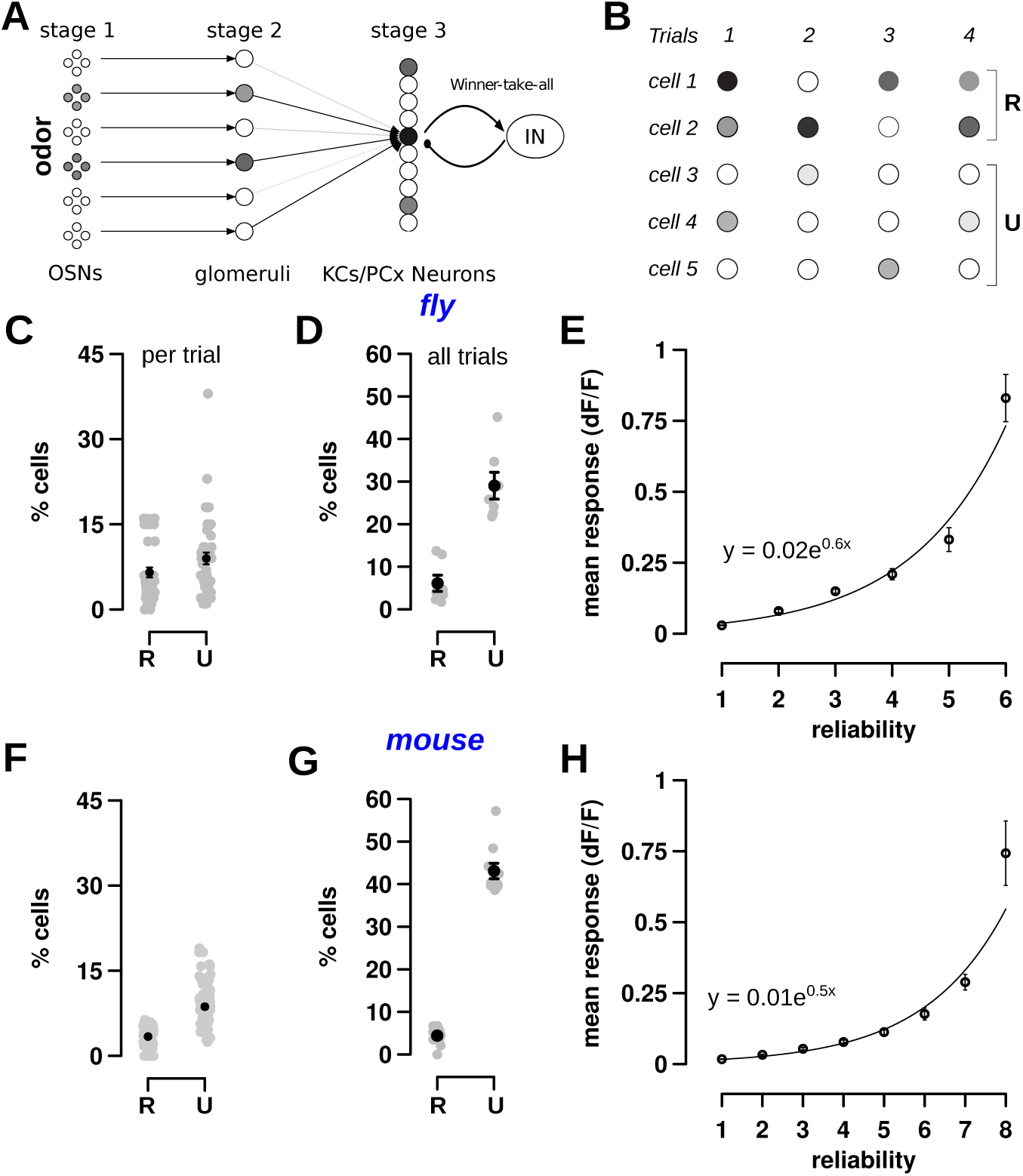
Response reliability is similar in third order neurons of the fly and mouse olfactory circuit. (A) Schematic of the first 3 stages of fly and mouse olfactory circuits. In stage 1, odors are sensed by olfactory sensory neurons (OSNs). In stage 2, OSNs pass odor information onto glomeruli structures in the antenna lobe (flies) or olfactory bulb (mice). In stage 3, odor information is passed to a larger number of Kenyon cells in the Mushroom Body (flies) or cells in the Piriform Cortex (mouse). Stage 3 cells are sparsified by an inhibitory winner-take-all circuit, which suppresses the activity of less responsive cells. (B) An example schematic showing response variability in stage 3 cells. Some cells (cells 1 and 2) have a large response in most trials (indicated by R for reliable cells), while most cells respond in *≤* half the trials with a smaller response (indicated by U for unreliable cells). (C–E) Responses of 124 Kenyon cells in flies for 7 odors with 6 trials per odor; data from Campbell et al. (2013). (C) The percentage of responsive KCs that are reliable (5.26 *±* 0.7%) and unreliable (7.23 *±* 0.84%) per odor trial. There are 42 grey points, one per odor-trial pair. (D) The percentage of KCs that are reliable (6.1 *±* 1.9%) and unreliable (29.0 *±* 3.16%) across all trials. Each grey point is an odor and the black point represents the mean. There are many more unreliable cells than reliable cells. (E) The mean response of a Kenyon cell (y-axis) increases with the cell’s reliability (x-axis). The measure of the fit, using linear regression on a log-plot, is shown in Figure S1A (*r* = 0.97). (F–H) Responses of 285 piriform cortex cells in mice for 10 odors with 8 trials per odor; data collected in this study. (F) The percentage of responsive PCx cells that are reliable (3.3 *±* 0.2%) and unreliable (8.7 *±* 0.4%) per trial. There are 80 grey points, one per odor-trial pair. (G) The percentage of PCx cells that are reliable (4.4 *±* 0.7%) and unreliable (43.1 *±* 1.8%) across all trials. Again, there are many more unreliable cells than reliable cells. (H) The mean response of a PCx cell (y-axis) increases with the cell’s reliability (x-axis). The measure of the fit is shown in Figure S1B (*r* = 0.98). Error bars in all panels indicate mean *±* SEM.

Here, we examined the sparsity and variability of responses to odors in the third stage of the circuit.

### Reliable and unreliable components of odor representations in fly mushroom body and mouse piriform cortex

**Fly.** The original studies of population activity in KCs measured sparseness using a particular response criteria (Campbell et al., 2013). They had shown that while odor responses in the Mushroom Body (MB) consistently evoked activity in about 5% of Kenyon cells across trials, the number of cells active per trial was much higher at about 12% (Methods). Here, we used those same criteria to group cells into reliable (the consistent 5% cells) and unreliable (the other) cells as illustrated in Figure 1B. *Reliable* cells are those that respond in more than half the trials (e.g., cell 1 with a reliability of 3 out of 4 trials, where reliability is the number of responsive trials), and *unreliable* cells are those that respond in *≤* half of the trials (e.g., cell 3 with a reliability of 1). We applied these criteria to the dataset — containing responses of 124 Kenyon cells in one fly to 7 odors with 6 trials per odor — to investigate the sparsity and variability of odor responses in the MB. This experiment was repeated in 5 other flies with similar results (Methods, Table S3).

Figure 1B illustrates how the responses of KCs vary across trials. On an average trial, about 13% of the 124 KCs exhibited a significant response, with 5% of those cells being reliable, and 7% being unreliable (Fig. 1C). By definition, the subset of reliable cells will mostly be the same across trials, whereas unreliable cells in one trial may differ from unreliable cells of other trials. Indeed, when examined over all trials (Fig. 1D), a total of 29% of the 124 KCs were unreliable, whereas 6% of the 124 KCs were reliable, which, as expected, is similar to the 5% fraction on an average trial. Figure 1B provides a visual illustration of this reliable-unreliable cell dichotomy. There is only one unreliable cell per trial (e.g., cell 4 in trial 1), but there are a total of three unreliable cells across all trials (cells 3–5). Thus, on an average single trial, 1 out of 5 cells (20%) are unreliable, and across all trials, 3 out of 5 cells (60%) are unreliable. On the other hand, on an average single trial, 1.5 cells are reliable, and across all trials, 2 out of 5 cells are reliable.

Reliable and unreliable cells also differed in the amplitude of their responses. As the reliability of individual KCs increased, their mean response levels increased exponentially (Fig. 1E; see Fig. S1A for fit of the linearized form, Pearson’s *r* = 0.97). While Figure 1E shows the mean for all KCs with the same reliability, Figure S1C shows each KC plotted individually and demonstrates a similar trend.

Thus, average measures of activity do not capture the diversity of cellular response properties in third order olfactory neurons. Responses on individual odor trials are composed of a core set of reliable KCs (about 5% of the population) as well as a peripheral cohort of unreliable cells (about 7% of the population in a single trial; 29% of the population across trials). We next asked if a similar stochastic code is observed in the mouse analog of MB: the piriform cortex.

**Mouse.** We collected new mouse data by imaging responses of 285 PCx cells to a diverse set of 10 odors with 8 trials per odor (Methods).

The responses of PCx cells bore close resemblance to KC responses in three ways. First, a small fraction (3.3%) of PCx cells responded reliably in each odor trial, with a larger fraction (9%) of unreliable cells (Fig. 1F). Second, the percentage of reliable cells across all trials was 4.4%, which is close to the percent of reliable cells in an average single trial; in contrast, the percentage of unreliable cells across all trials was much higher: 43% (Fig. 1G). It is possible, as we show later, that the increase in the number of unreliable cells in the mouse compared to the fly is due to the increase in the number of trials analyzed (6 trials per odor in the fly versus 8 trials per odor in the mouse). Third, as the number of trials in which a PCx cell responds increases, the size of its response increased exponentially (Fig. 1H; Fig. S1B shows linearized fit, *r* = 0.98). These results were repeatable for 4 other mice (Methods, Table S4).

Thus, odor responses of cells in MB and PCx are similar with respect to the parameters analyzed. The responsive population contains a small number of cells that respond reliably across trials with large responses, while a larger number of cells respond unreliably or infrequently with smaller responses.

### Cell reliability, response size, and odor specificity levels lie on a continuum

Thus far, we classified a cell based on whether it responded in at least one and up to half of the trials (unreliable) or if it responded in more than half of the trials (reliable). This division into two classes was used to ease exposition, however, it is somewhat arbitrary. Indeed, there are gradual differences in response sizes between cells based on their reliability (Fig. 1E,H). We next sought to determine whether three properties of cells — cell reliability, response size, and odor overlap (described below) (Fig. 2A) — fall on a continuum, and if there is some structure to this distribution.

**Figure 2:**
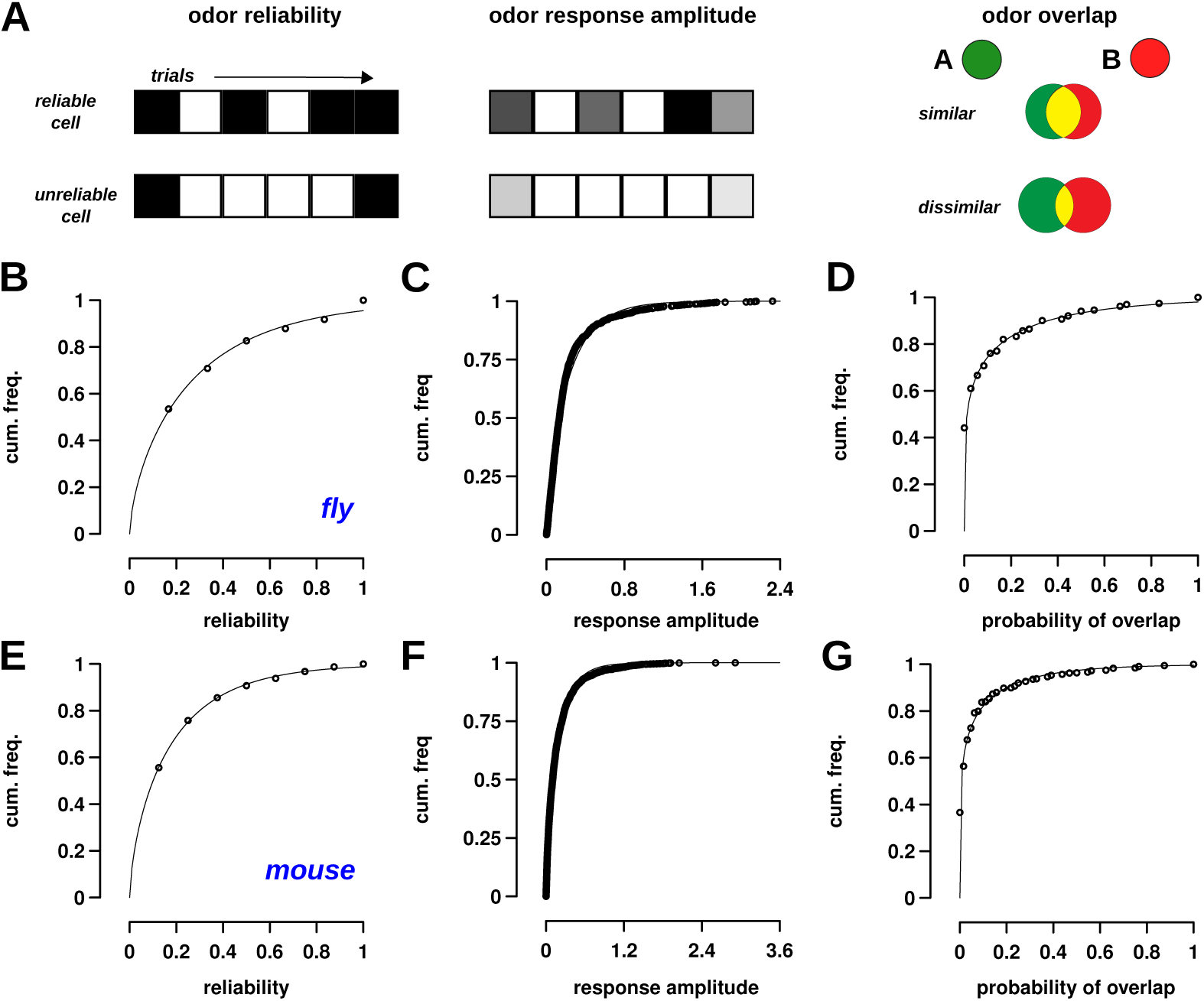
Cell reliability, response size, and odor specificity follow a continuous distribution. (A) Schematic of three response properties that were examined for odors and odor-pairs. Odor reliability refers to the number of trials for which a cell is responsive. Each trial is depicted by a grid square, and black color indicates a response. Cells that respond in more than half the trials were classified as reliable; all other responsive cells were unreliable. Odor response size is the firing rate of the cell in individual trials, as measured by Ca^2+^ flourescence. The top line shows a cell with a large response, where the intensity of grey denotes size of response. Odor overlap is the probability that the cell will respond to both odors. The odor overlap here is shown in yellow and decreases for dissimilar odors (bottom) compared to similar odors (top). (B,E) Cumulative frequency histograms showing the distribution of reliabilities for Kenyon cells (flies) and piriform cortex cells (mice). Each circle represents the cumulative probability for cells with a reliability value represented on the x-axis. The points are fit by Gamma distributions with shape = 0.64, scale = 0.42 (for B, fly), and shape = 0.64, scale = 0.17 (for E, mice). The significance of the Gamma distribution is that it is a maximum entropy code and optimizes for the most stimuli that can be encoded (see text). (C,F) Cumulative frequency histograms for response sizes in both flies and mice are well fit by Gamma distributions with shape = 0.77, scale = 0.28 (for C, fly), and shape = 0.70, scale = 0.24 (for F, mice). (D,G) Cumulative frequency histograms for the overlap of cells between odor-pairs in both flies and mice, also fit using Gamma distributions with shape = 0.19, scale = 0.60 (for D, fly), and shape = 0.18, scale = 0.35 (for G, mice).

The distribution of reliability values of KCs and PCx cells was well-fit to a Gamma distribution in both flies (Fig. 2B) and mice (Fig. 2E). The parameters of the two Gamma distributions were almost identical; fly: shape=0.64, scale=0.42; mice: shape=0.64, scale=0.28. Most cells have a low reliability — in Figure 2B, the lowest reliability plotted (0.167) starts at a cumulative frequency of 0.5 — whereas a few cells have high reliability (the upper part of the curve), reflecting our previous results that there are more unreliable cells than reliable cells. Thus, reliability is a property that changes gradually and is not bimodal with two mutually exclusive classes.

The distribution of response sizes of KCs and PCx cells also fit closely to a Gamma distribution in both flies (Fig. 2C) and mice (Fig. 2F). Again, the parameters of the Gamma distributions were similar; fly: shape=0.77, scale=0.28; mouse: shape=0.70, scale=0.24. This result also confirms previous studies by (Stevens, 2015, 2016), who showed that olfactory neuron responses follow a maximum entropy code (Jaynes, 1957). A code is called maximum entropy if it can encode the maximum number of stimuli using a given population of neurons and set of constraints. Without any knowledge of the statistics of the population responses, the maximum entropy code is a uniform distribution. For instance, consider a neuron with a firing rate range of 0–100 spikes/s, and it has to encode odor concentration in the range of 0–100 ppm. The neuron’s coding would be inefficient if concentrations 0–80 ppm were represented by firing in the range of 0–50 spikes/s. The most efficient coding system would be to uniformly map concentration to response levels (0–100 ppm to 0–100 spikes/s) (Laughlin, 1981). If, however, the mean of the population is constrained (mean response rate of an odor across all cells or of a cell across all odors is the same), the maximum entropy code follows an exponential distribution of firing rates. Analysis of responses in the first two stages of the fly circuit (Stevens, 2015, 2016) showed that each odor’s response follows an exponential distribution such that only a few cells are highly active for each odor, and the mean activity rates of different cells was the same (constrained mean). The study predicted that the distribution of responses in the third stage (Kenyon cells) would be a Gamma distribution, because each KC integrates 6-8 PN inputs, and a convolution of exponentials is a Gamma distribution. Incidentally, a Gamma distribution is also a maximum entropy code, wherein the mean and log(mean) are constrained. Our results show that MB and PCx responses indeed follow a Gamma distribution. Both regions are optimized to encode the maximum number of stimuli. Thus, reliability and response level properties of cells in MB and PCx change gradually.

We developed a third property, which we call *overlap*, to capture the degree of similarity in cells’ responses to pairs of odors. First, for each cell, we computed its response probability (i.e., whether it will respond to the odor in a randomly chosen trial); e.g., if a cell responds in 4 out of 6 trials, its response probability is 4/6 or 2/3. This measure is also the reliability measure of the cell used in previous figures (Figs. 1,2). Next, we calculated the probability that the cell will respond to both odors, which is the product of response probabilities to the two odors. For instance, if a cell responds to the first odor in 4 out of 6 trials, and to the second odor in 2 out of 6 trials, the probability of responding to both odors is 8/36, and the overlap score is 0.22 (Fig. S4C for an example odor-pair). We repeated this procedure for every responsive cell across all odor-pairs. Then, we calculated the mean overlap value across all cells for each pair of odors. We found that the distribution of overlap values of KCs and PCx cells to pairs of odors also followed a Gamma distribution in both flies (Fig. 2D) and mice (Fig. 2G), with similar parameters: fly: shape=0.19, scale=0.6; mouse: shape=0.18, scale=0.35 (Fig. S4A,B for all odor-pair points plotted by reliability). The distribution implies that 80% of cells have overlap *<*0.2 and that 40% of cells are grouped at the lowest overlap value. Bias in odor selection is unlikely to drive these relationships since the similarity between the pairs of odors (measured as the correlation between KC representations of odors) is uniformly distributed for flies and mice (Fig. S5) between 0 and 1. Thus, Figure 2D,G shows that most cells have very little similarity across odor-pairs in terms of their response. It raises two questions relevant to discrimination: do cells with high overlap belong to similar odors, and are low overlap cells also more unreliable? (as Fig. 1 showed unreliable cells differ even in between trials of the same odor). We tackle both questions and their implications for discrimination in subsequent sections (Figs. 4,5,6).

**Figure 3:**
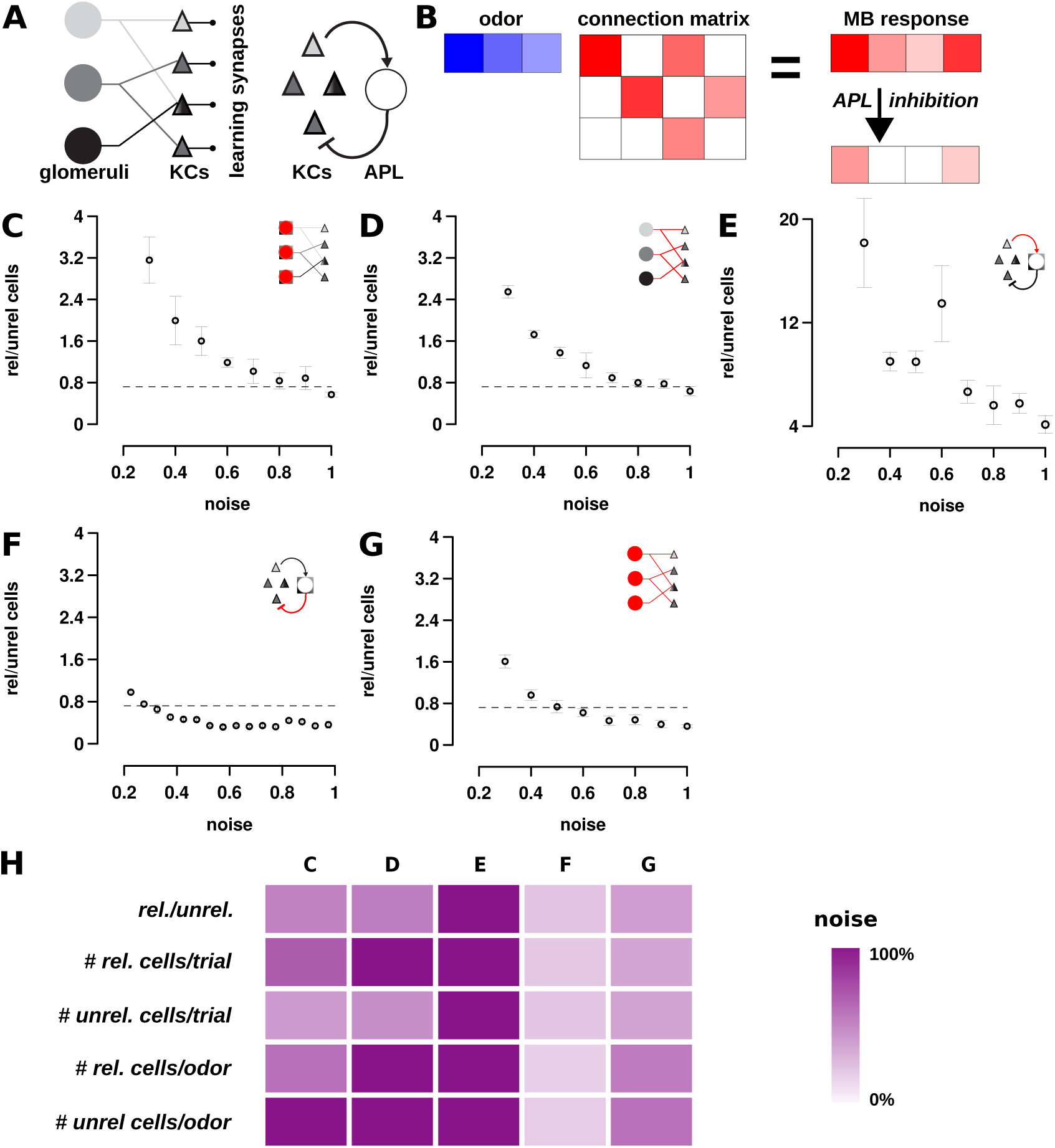
A winner-take-all mechanism is required for generating stochastic codes. (A, left) Schematic of information transfer in the fly olfactory circuit. Odor information from OSNs is passed from glomeruli to Kenyon cells (via projection neurons, PNs) in the mushroom body. Kenyon cells synapse with MBONs that influence behavior. The KC-MBON synapses are subject to synaptic plasticity. (A, right) The winner-take-all (WTA) circuit in MB, where all KCs activate the inhibitory Anterior Paired Lateral neuron that in turn feeds back and inhibits all KCs. (B) A schematic of the linear rate firing model. The PN response to odors is depicted in blue, and their connection matrix with KCs is denoted in red. With a linear rate firing model, we take a product of the odor vector and connection matrix to get the KC response, which is sparsified by the APL neuron inhibitory feedback. (C–G) Model simulations show that injecting noise in different parts of the circuit (C) PNs, (D) PN*→*KC synapses, (E) KC-APL synapses (F) APL-KC synapses, and (G) PNs and PN*→*KC synapses produces different ratios of reliable to unreliable cells responding per trial. The straight line in the plots denotes reliable/unreliable cell ratio of 0.72 observed with experimental MB responses (Fig. 1). Model (F) requires the least amount of noise (intersection of line and data points at x = 0.275 in F) to observe this ratio. For all plots, the x-axis denotes the fraction of injected noise. Thus, 1 denotes that noise levels are 100% of signal. We show noise levels up to 100%, as the noise levels above it do not produce desired outcomes. (H) Summary of how well each model recapitulated the KC response statistics observed in Figure 1. Each row denotes a response statistic, e.g., the reliable/unreliable ratio. The color intensity indicates the amount of noise required to get response statistics that match data. Darker colors indicate lower amounts of noise. Adding noise to the APL-KC connections produces observed responses with the least amount of noise. For combination model G, all components are perturbed with the same amount of noise; for effects of varying amounts of noise, see Table S1 and Methods.

**Figure 4:**
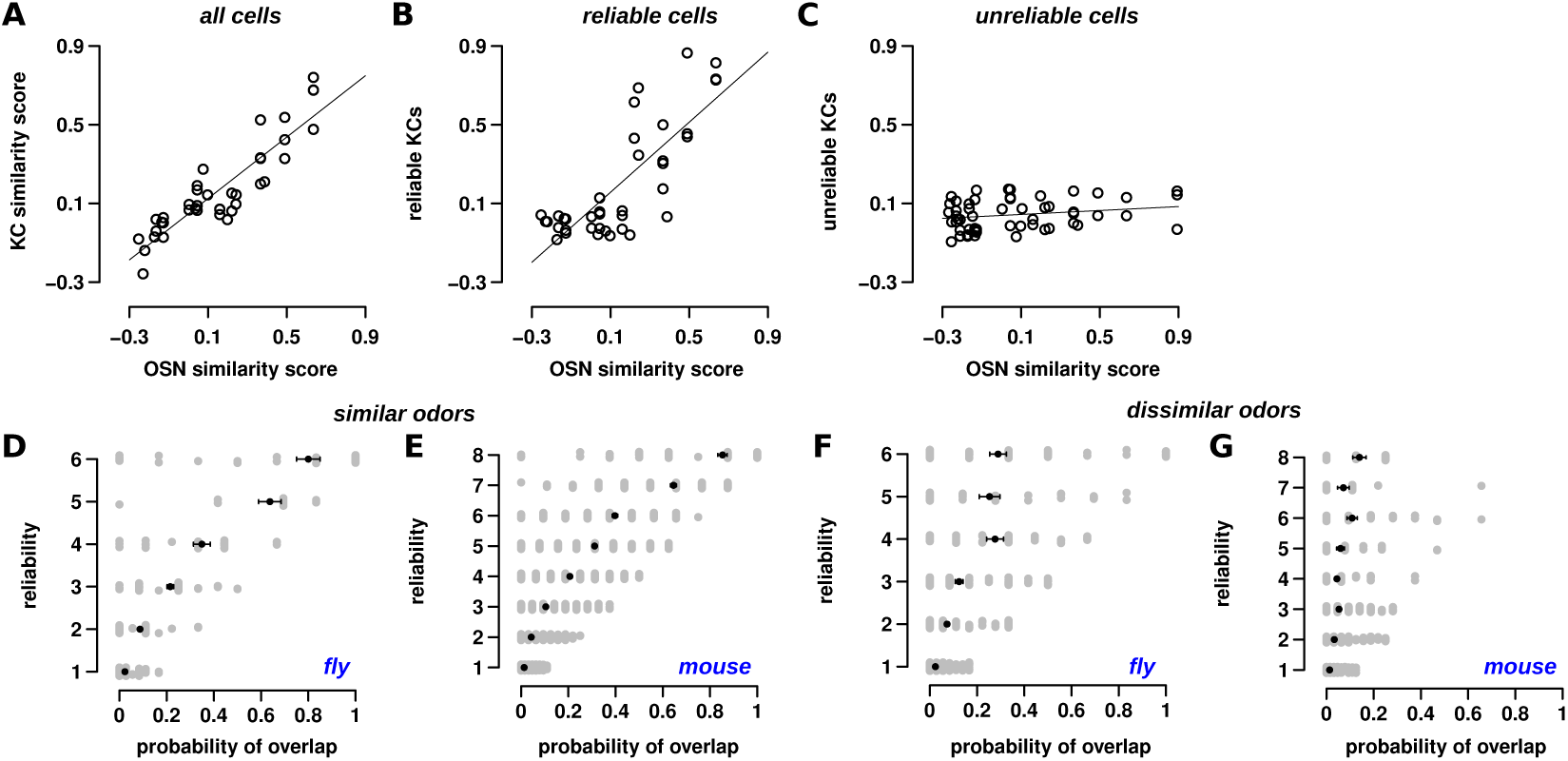
Reliable and unreliable cells differ in their response to odors. (A) The similarity between odor pairs was calculated based on OSN response correlation in the fly’s antenna (Hallem et al., 2006) (x-axis) and KC response correlation in the fly’s mushroom body (y-axis). Each circle denotes one odor-pair. Similarity between odor pairs was calculated via Pearson’s correlation of response vectors. When considering all KCs, odor similarity was highly correlated (*r* = 0.89). The equation for the line fit is *y* = 0.78*x*. (B) When considering only reliable Kenyon cells, odor similarity is also highly correlated (*r* = 0.80). The equation for the line fit is *y* = 0.88*x*. (C) In contrast, the similarity between odor-pair responses for only unreliable cells has very low correlation (*r* = 0.20). The line fit has equation *y* = 0.05*x*, i.e., the slope is close to 0. In other words, odor pairs that are dissimilar (left part of the x-axis) have only slightly less correlation than similar odor pairs (right part of the x-axis). (D,E) In both flies and mice, reliable cells overlap more than unreliable cells for similar odors (i.e., odor-pairs with correlation in their odor responses *>* 0.5). Here, overlap is the probability that a cell will respond to both odors, e.g., if the probability of response to the two odors is 1/6 and 3/6, the overlap is 3/36 or 1/12. Each grey circle shows the overlap between two odors for a single cell, whose reliability for the first of the odor pair is shown on the y-axis. Pearson’s correlation between overlap and reliability is 0.87 and 0.91, for flies and mouse, respectively. (F,G) For dissimilar odors (correlation *<* 0.15), while more reliable cells have a higher overlap than unreliable cells, the increase in overlap is much less than for similar odors shown in (D,E). Correlation is 0.55 and 0.33 for linear fits, with slopes of 0.05 and 0.009, respectively, for flies and mice.

**Figure 5:**
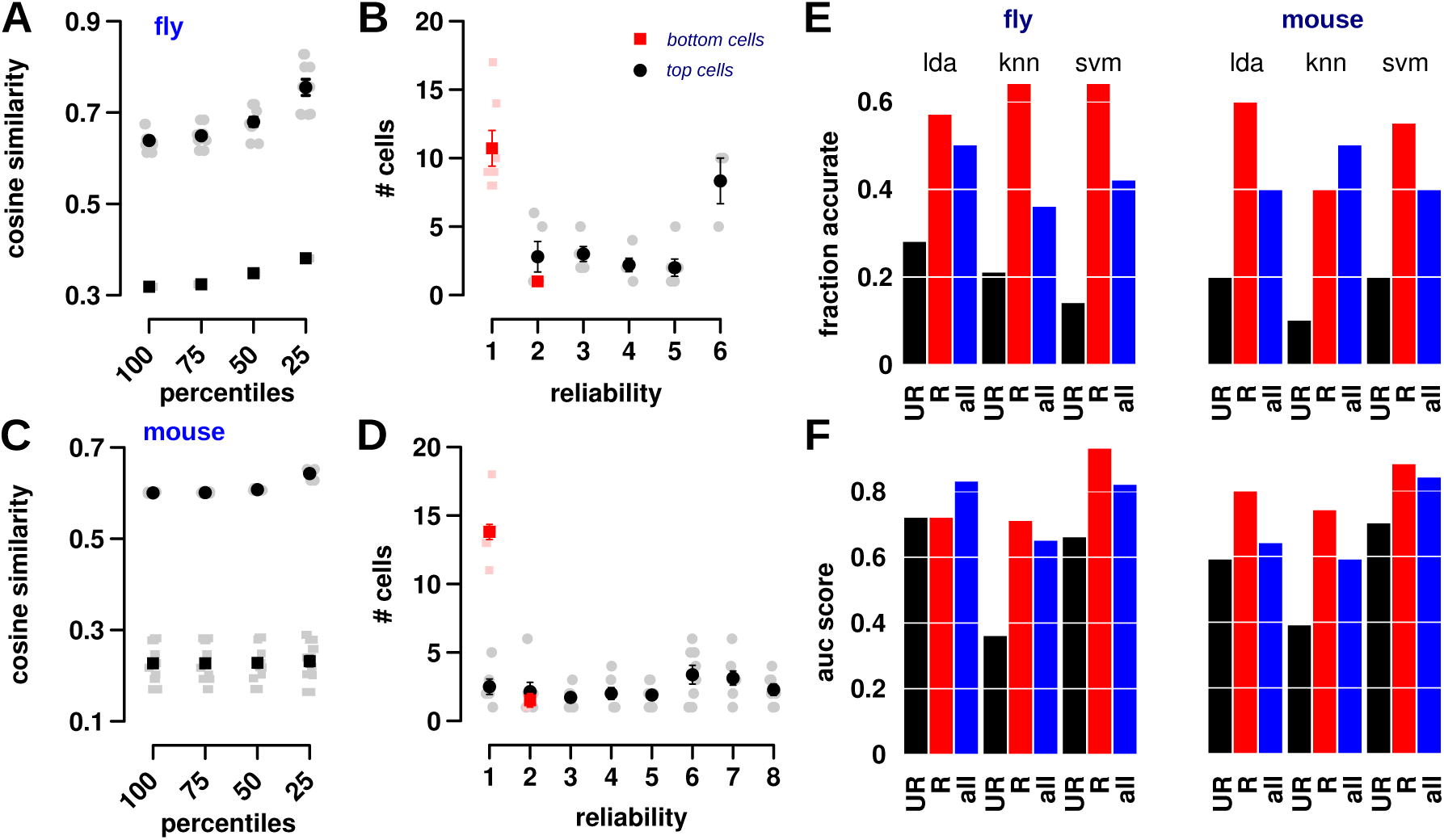
Reliable and unreliable cells contribute to odor discrimination. (A) The cosine similarity (y-axis) for the top 25, 50, 75, and 100% of responsive Kenyon cells (x-axis) for similar odors (correlation *>* 0.5, top circles) and dissimilar odors (correlation *≤* 0.15, bottom squares). Each square and circle here are odor pairs. The top 25% of cells are more similar than the rest of the population (i.e., they provide no extra discriminatory information). The cosine similarity drops significantly for dissimilar odors, suggesting that there is significant discriminatory information, regardless of what percent of the responsive population is analyzed. (B) The number of cells at each reliability level for the top 25% (black circle) and bottom 25% (red square) of cells. Most of the top 25% of cells are reliable, while nearly all of the bottom 25% of cells have the lowest reliability. (C) The same plot for mice as (A), showing similar trends. (D) The same plot for mice as (B), showing similar trends. (E) Simple machine learning algorithms applied to reliable, unreliable, and all cells show that reliable cells are significantly better at classification than unreliable cells. Compare the length of the red bars (reliable, R) to the black bars (unreliable, UR), where the length of the bar denotes the fraction of correct classifications. The performance of decoders with all cells (blue bars) is similar to reliable cells, though slightly less. We used three decoders (svm – support vector machines, knn – k nearest neighbor, and lda – linear discriminant analysis). See Methods and Table S2 for algorithm details and parameters for the decoder, and training and test sets. (F) Area under the curve measure (averaged over all pairs of odors) for assessing the pair-wise discrimination performance of unreliable (U, black bars), reliable (R, red bars), and all (blue bars) cells for the three decoders (lda, knn, and svm). Reliable cells, on average, have slightly higher AUC measures. Unlike accuracy measures from (D), however, unreliable cells (in black) show significantly higher AUC measures, suggesting that they provide significant information for distinguishing odors. See Figure S3D for area under the ROC (AUC) measure for assessing performance of the classifiers specifically for similar and dissimilar odors.

**Figure 6:**
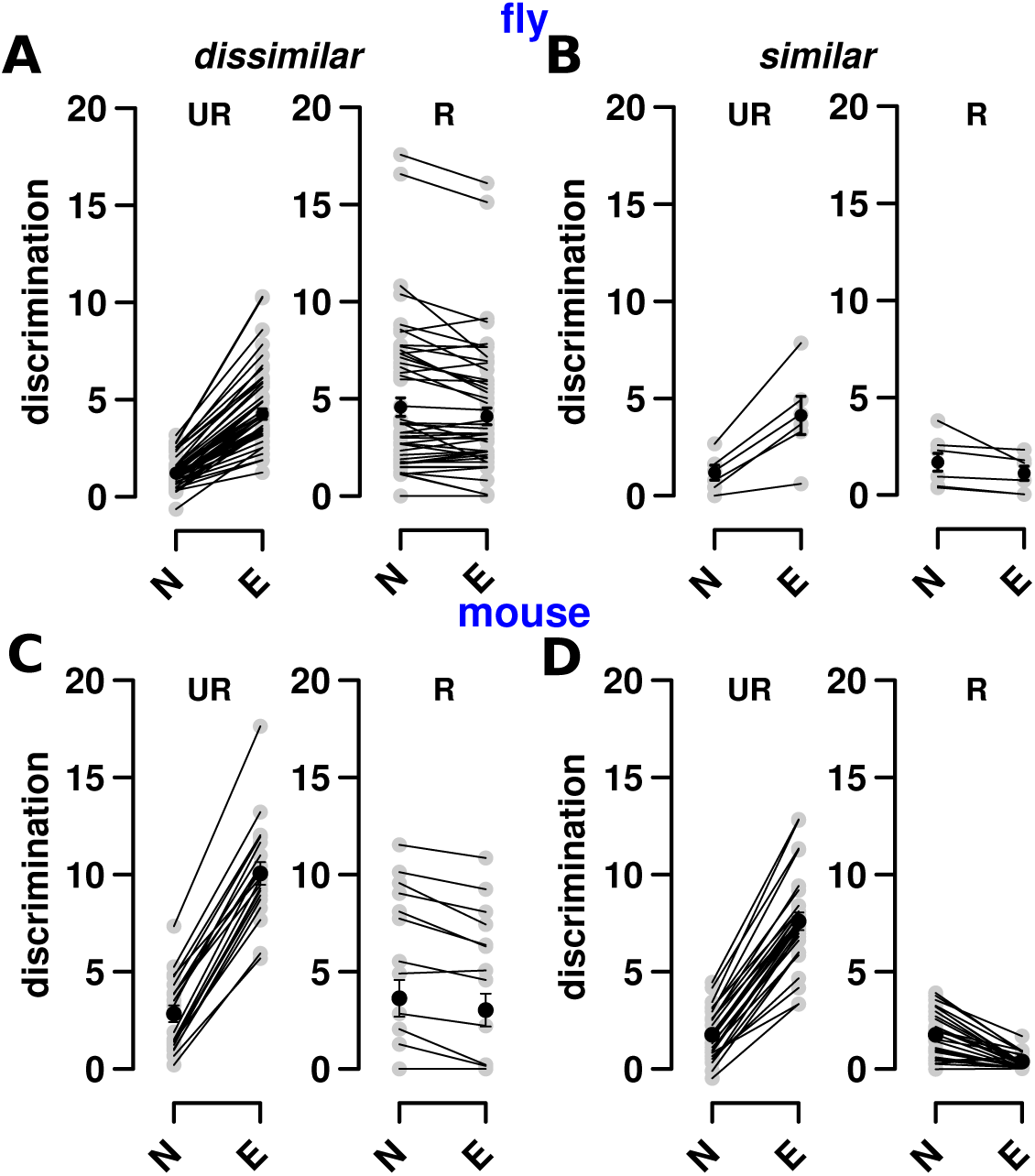
Unreliable cells contribute more to discrimination of similar odors. Extended learning increases the amount of discriminatory power between odor-pairs. Odor-pairs are split into dissimilar (A,C) odor pairs, whose correlation coefficient is less than 0.15, and similar (B,D) odor-pairs with a correlation greater than 0.5. (A) Shown are two plots of how extended (E, x-axis) and normal (N, x-axis) training affect the contributions of (y-axis) unreliable cells (left plot, UR) and reliable cells (right plot, R) towards discrimination of dissimilar odors. Each grey point represents the discrimination measure (eq. 2) for an odor-pair. Discrimination calculations for the fly are based on data from Campbell et al. (2013). Unreliable cell contributions increase with extended training, to match the contributions of reliable cells. (B) Similar plots as (A) for similar odors. Here, too, extended training increases the discrimination contribution of unreliable (UR) cells. For reliable cells, however, contributions are maintained or reduced. (C-D) are equivalent calculations for the mouse data set shown in Figure 1.

These results highlight two characteristics of olfactory responses. First, every odor-cell pair is defined by its response level and reliability. Both characteristics are tightly coupled, such that if a cell has a large response to an odor, it is also highly likely to respond in most trials. Second, all three properties follow a Gamma distribution, which indicates that cells do not fall into two simple categories, but rather along a continuum.

The large number of unreliable cells raises the issue that these cells, although significantly responsive for a trial, may respond only in that trial and then stay silent. In such a case, even if these cells are active during training, they will not influence subsequent behaviors. We found, however, that this is unlikely. We re-examined fly and mouse datasets, except this time we considered only the first 3 trials, and isolated those cells that responded only once — these would constitute the unreliable cells if the experiments included just the first three trials (Fig. S6A–C). We then calculated their response frequency in the remaining trials (second 3 trials for flies, and trials 4–8 for mice), and found that responses did not match the predicted frequency of 1/3 (Fig. S6D–F). Instead, the responsive population is a composite of cells with a range of reliabilities, showing that even unreliable cells are likely to respond again, though with varying length of intervals in between, consistent with cell reliabilities following a continuous distribution (Fig. 2 shown above). Thus, even cells with low reliability can play a role in subsequent odor learning and behavior.

### Circuit mechanism for producing reliable and unreliable cells

Where does response variability come from? Variability is typically ascribed to sensory noise or noise in inputs to a brain region (like neuromodulators, for instance). These inputs to MB are known (Li et al., 2020; Takemura et al., 2017), and thus, MB presents an excellent substrate to enquire whether experimental or sensory noise generates the response variability observed, or if other mechanisms are involved.

To explore how noise introduced at different stages in the circuit (Fig. 1A) affects the emergence of reliable and unreliable cells, we developed a linear rate firing model of the fly olfactory circuit (Fig. 3A,B). In a linear rate firing model, we treat the inner workings of each neuron as a black box and assume that its output is a linear function of its inputs. Although, it does not capture a neuron’s inner workings, it does well in recapitulating the dynamics of the olfactory circuit network as a whole in addition to simplifying analysis as successful approaches in the past have demonstrated (Luo et al., 2010; Barak et al., 2013; Stevens, 2016). As described in Methods, the PN*→*KC connection structure was based on Caron et al. (2013) and Gruntman & Turner (2013) and subsequent analysis of this data by Stevens (2015); specifically, each of the 2000 KCs receives synapses from approximately 6–8 of the 50 PN types, with synapse strengths following a Gamma distribution. The distributions of KC*→*APL and APL*→*KC synapse strengths were based on Li et al. (2020) and Takemura et al. (2017) (see Methods for statistics).

We introduced noise in firing rates of cells (PNs) and in synaptic transmission between PN*→*KCs, KCs*→*APL, and APL*→*KCs. Thus, we tested four manipulations. For each manipulation, we added multiplicative noise by sampling from a Gaussian distribution with a mean of 0 and an SD in the range of 0–150%. For instance, to add 15% firing rate noise to a PN with response amplitude, *P*_resp_, we first computed the noise as *P_η_* = *N* (0, 0.15), and then computed the noisy response as *P*_resp_(1 +*P_η_*). The noise component for each response was generated afresh for each trial. Our goal was to determine which of the four manipulations, under “reasonable levels” of noise, could generate a stochastic code (i.e., reliable and unreliable cells for an odor). These noise additions could arise here, as it does in many biological systems (Hornung & Barkai, 2008; Sterling & Laughlin, 2015), through variations in component number within the transmission machinery such as neurotransmitters or synaptic vesicles or vesicle release probability (Franks et al., 2003; Bekkers et al., 1990; Faisal et al., 2008). A reasonable level of noise for PNs would be in the range 0–50% (Bhandawat et al., 2007), and for synaptic transmission, would be in the range of 15-25% (Bekkers et al., 1990; Franks et al., 2003).

We reasoned that a plausible noise source should be able to replicate the following five characteristics of the olfactory stochastic code shown in Figure 1: i) a ratio of reliable to unreliable cells of 0.72, ii) # reliable cells per trial of 5.3, iii) unreliable cells/trial = 7.2, iv) reliable cells/odor = 6.1, v) unreliable cells/odor = 29. When physiologically reasonable levels of noise was introduced in the APL*→*KC synapse, the five characteristics were well-matched with observations (Fig. 3). Other components of the circuit either required unrealistic levels of noise (around 80–100% for PN and PN*→*KC noise, Fig. 3C,D) or did not match the reliable/unreliable cell ratio of 0.72 under any level of noise tested (KC*→*APL noise, Fig. 3E).

We also examined how multiple noise sources could interact. We exhaustively searched parameter space by combing through all combinations of PN, PN*→*KC, APL*→*KC noise levels, and APL*↔*KC gain levels to isolate those noise combinations that produced all 5 characteristics of the code. See Table S1 and Methods:Noise exploration section for a detailed description of parameter explorations. Noise in the range of 0-100% was added to all components of the model, and we selected combinations that satisfied all 5 constrains (Fig. 1, listed above). Of the 44,760 combinations, only 2,500 satisfied this criterion, and the top 700 of them had noise in APL*→*KC connections in the 15–30% range. Noise in the other components by themselves did not satisfy all constraints, showing that APL*→*KC noise is essential to generating the stochastic code.

Further analysis also revealed how noise in the winner–take–all (WTA) circuit produced a stochastic code. We used data from Lin et al. (2014), who silenced APL and recorded Ca^2+^ responses from KCs (to three odors in this analysis and across 10 trials in different flies). KC odor responses without APL was well-fit by a Gamma distribution with parameters of shape= 4.2, and scale = 5.8 (Fig. S7B–D). We then introduced WTA feedback such that all but the top 10% of cells were silenced. When we added 15–20% noise in the APL to KC feedback, we observed a stochastic response with 3 properties. First, KCs with response levels close to the threshold percentile (in this case 90%), showed the most unreliable responses across trials; i.e., they were the most sensitive to noise. Second, KCs that had a higher percentile score were more reliable; e.g., KCs that had a percentile score of 98% were more likely to respond in nearly all the trials, much like reliable cells. Third, conversely, KCs that had a lower percentile score were less likely to respond (i.e., stay silent) as more noise was needed to push them past the threshold, and since high amplitude noise events are less likely, the cells respond less.

The WTA model recapitulated two important characteristics of the data. First, as Kenyon cell reliabilities increased, the average response amplitude also increased. Second, while the number of reliable cells was around 6%, the number of cells responding in each trial was close to 14%, and in total, across 6 trials, there were 25–35% unreliable cells, much like what we observed with the data (Fig. 1). In conclusion, our model shows that the WTA mechanism is necessary and sufficient for producing observed results (although some sensory noise still exists), suggesting a mechanism for generating reliable and unreliable cells.

### Reliable and unreliable cells encode different amounts of odor information

#### Preservation of odor similarity from Circuit level 1 to Circuit level 3 olfactory cells

To gain insight into the functional differences of the two classes of cells, we next asked: Are reliable or unreliable cells better at preserving odor similarity as it transfers from the antennal lobe to Mushroom Body (MB)? To answer this question, we compared the similarity of population response patterns in OSNs with the similarity of KC responses across a series of odor pairs. We then split KCs into reliable and unreliable classes and tested how well each class preserved odor similarity. For OSN responses, we used the dataset from Hallem & Carlson (2006), who recorded responses of 24 OSN types in the antenna lobe to the same set of odors that were used by Campbell et al. (2013). For MB responses, we used the set of 124 KC responses provided by Campbell et al. (2013).

Consistent with Schaffer et al. (2018), we found that most of the odor information in the OSN population is preserved in MB; the Pearson correlation coefficient of odor-pair responses in OSN and KC populations is *r* = 0.89, with a slope of 0.78 (Fig. 4A). The slope captures how well the relationship between similar and dissimilar odors is maintained from the antenna to MB; a slope of 1 means perfect maintenance.

When splitting KCs into two classes, we found that reliable KCs better preserved the similarity of OSN representations and had a higher slope (*r* = 0.80, slope=0.88; Fig. 4B) compared to unreliable KCs (*r* = 0.20, slope=0.05; Fig. 4C). When odors have dissimilar OSN representations (low correlation), reliable KC population responses also have low correlation, and the reliable KC correlation increases with OSN represention similarity. For unreliable cells, the correlation trend is markedly different, with a low slope and correlation when comparing the similarities between OSN and KC representations. When odors have dissimilar OSN representations, unreliable KC populations have low correlation just like their reliable counterparts. But unlike reliable KCs, as OSN representation similarity increases, there is only a slight increase in unreliable KC correlation, showing that unreliable cells are decorrelated whether odors are similar or dissimilar.

We could not carry out a similar analysis with the mouse dataset, as we did not have odor responses to the same odors in the olfactory epithelium and PCx. We, therefore, compared the similarity of odor-pairs with all PCx cells versus just the cohort of reliable or unreliable cells (Fig. S3E,F). First, however, we carried out the comparison for KCs, and found that this analysis recapitulated the results of Figure 4A-C: reliable cell and “all cell” population similarities (of odor–pairs) were proportional while those of “all cells” and unreliable cells were decorrelated. Repeating the same analysis on PCx cells revealed a similar trend, suggesting that in mouse PCx cells, too, unreliable cells are decorrelated between all odor-pairs, while reliable cells are decorrelated only amongst dissimilr odors (with low correlation between odor–pairs).

Next, as alternative analysis, we used the overlap measure, introduced earlier (Fig. 2D,G), to capture how reliable and unreliable cells responses change with odor similarity. For each set of cells (reliable or unreliable), we calculated the individual overlap measure of each cell and averaged the whole population. Thus, this overlap measure computes how well reliable and unreliable cell populations preserve odor similarity. We defined similar odors as those odor pairs with a correlation *>* 0.5 in both MB and PCx (4 similar odor-pairs in flies and 3 similar odor-pairs in mice). Conversely, odor-pairs that had a correlation *<* 0.15 were designated as dissimilar odors (8 dissimilar odor-pairs in flies and 7 dissimilar odor-pairs in mice).

For similar odors, in both flies and mice, the higher the reliability of a cell-odor pair (the average reliability of a cell to both odors), the higher the overlap (Figs. 4E–F). In other words, as overlap increases (*x*-axis in Figs. 4E–F), the reliability (*y*axis) increases proportionally: fly: *r* = 0.87, mouse: *r* = 0.91. Conversely, for dissimilar odor pairs (Figs. 4G–H), as overlap increases, reliability increases more gradually: fly: *r* = 0.55, mouse: *r* = 0.33. As above, the slopes of the correlation lines also slightly decreased from reliable cells (fly: 0.15, mouse: 0.10) to unreliable cells (fly: 0.05, mouse: 0.009). The slope indicates the relationship between overlap and reliability: the higher the slope, the higher the dependence between the two. Thus, when odors are similar, reliable cells are likely to respond more similarly than their unreliable cells.

### Reliable cells cannot by themselves distinguish similar odors

Studies have suggested that sparse coding, which restricts odor coding to the top firing neurons, might aid in the discrimination of similar odor-pairs by reducing the overlap in their representations. Our findings suggest, however, that sparse coding might not reduce overlap, since reliable cells, which tend to have a larger response (Fig. 1E,H), are highly correlated (Fig. 4) for similar odor-pairs. Upon examining the fly and mouse datasets, we found, as expected, that the more reliable a cell, the more likely it is in the top 25% of highest firing neurons (Fig. 5B,D). Similarly, the more unreliable a cell, the more likely it is in the bottom 25% (Fig. 5B,D). Notably, the top responding cells, in both flies and mice, reflect the same trend as reliable cells: as odor similarity increases, so does the similarity of the top responding cells (Fig. S8).

We delved deeper into the consequences for sparse coding by looking at the cosine similarity between odors (KC vectors), for four sub-populations — the top 25, 50, 75, and 100 percentiles of responding cells — to test how sparsity level affects the overlap in odor representations (Fig. 5A,C). These are percentiles of only responsive cells for the odor, which means about 30% of the population in flies and about 40% in mice, on average. Strikingly, we found that the cosine similarity of odors did not decrease from the top 50% of responsive cells to the top 25% of cells for similar odors (top circles), which would be expected if sparse coding improves discrimination. For flies and mice (Figs. 5A,C), the cosine similarity between dissimilar odors (bottom squares) is much less than similar odors. If the circuit were to increase sparseness by simply changing the threshold of responding cells (by increasing inhibition, for example), this would not actually improve discrimination for similar odors, because the cells that are left to respond are reliable cells, which have high overlap and correlation (Fig. 4).

The previous analysis compared the overlap between percentiles of responsive cells, but the conclusion is similar when comparing the overlap between quartiles of responsive cells (i.e., instead of the top 0–25% vs. 0–50% we compared 0–25% vs. 25–50%; Figure S8A). The top 25% of cells, similar to reliable cells in Figure 4, are the best indicators of odor similarity shown by the greater slope: they are more similar when odors are similar and dissimilar when odors are dissimilar. Figure S8B shows a similar result for the mouse dataset, suggesting that sparse coding does not aid in pairwise discrimination of similar odors.

#### Discriminating odors using reliable and unreliable cells

Which cells, reliable or unreliable, are more informative in discerning odors? To answer the question, we used two approaches.

In the first approach, we used simple machine learning classifiers on the fly and mouse datasets. The premise is that a decoder might approximate decision-making neurons downstream of MB or PCx that identify odors and tell them apart, and, thus, might serve as a proxy for determining the amount of discriminatory information provided by reliable and unreliable cells. Briefly (see Methods for details), for each dataset, we isolated the reliable cells by setting the responses of unreliable cells to 0. We then split all odor trials into training (*∼*80% of trials) and test (*∼*20% of trials) sets (Methods for details of trials cross-validation details etc.). There were a total of 7 odors and 42 trials for flies, and 10 odors and 80 trials for mice. We repeated the same procedure for unreliable cells and *all cells* together. We applied this procedure to three decoder models: Linear Discriminant Analysis (LDA), *k* Nearest Neighbors (kNN), and Support Vector Machines (SVM) with a linear kernel.

Figure 5E shows the test accuracy of each decoder in predicting the correct identity of the odor. Reliable cells (red bars; the bars represent the fraction of odors that were correctly classified, and the values are presented in Table S2) are much better at classification compared to unreliable cells (black bars). Classifying with *all cells* is significantly better than with unreliable cells, though not as good as reliable cells.

Figure 5F shows the area under the curve (AUC) results of each decoder for the test set when telling apart pairs of odors. AUC for an odor-pair measures the number of true versus false positives as per Hand & Till (2001). Given two odors A and B, AUC is the probability that a random trial of odor A has a lower chance of being classified as odor B, than as odor A. Thus, a score of 1 shows that A can be easily distinguished from B, while 0 indicates a high likelihood of confusing the two odors. In general, odor-pairs that had low AUC had high correlation of their KC representations (Fig. S3D similar, r *≥* 0.5, AUC: 0.026), and odor pairs with high AUC had lower correlation of KC representations (Fig. S3D dissimilar, r *<* 0.25, AUC: 0.95). Figure 5F shows that reliable cells still provide, on average, a high level of distinguishing information (plotted values are presented in Table S2). What is also evident, however, is that unreliable cells, too, provide significant distinguishing information between any pair of odors. Thus, our analysis of the data using linear decoders and statistical analysis shows that discrimination between odors becomes hard when their similarity increases, reflecting experimental findings of difficulties organisms face when discerning similar odors (Campbell et al., 2013; Lin et al., 2014; Chapuis & Wilson, 2012).

So, what value might unreliable cells provide for distinguishing odors? Figure 5 shows that in some cases, reliable cells alone perform better at discrimination than all cells; however, Figures 4E–F shows that for similar odors, unreliable cells are less overlapping than reliable cells. How might the circuit take advantage of this non-overlap provided by unreliable cells to better distinguish similar odors?

To understand the contribution of unreliable cells, we used a learning model similar to the fly learning and discrimination system (Aso et al., 2014b; Modi et al., 2020; Aso & Rubin, 2016) (Fig. 7A), since its architecture is better mapped than that of the mammalian system. Learning in the fly system occurs by coordination amongst three sets of neurons: KCs that encode odor information, dopaminergic neurons or DANs that convey reinforcement signals (i.e., reward or punishment), and MB output neurons or MBONs that drive approach or avoidance behavior (Modi et al., 2020). During appetitive training, DANs for reward depress synapses between KCs active for the odor (conditioned stimulus or CS) and MBONs that drive avoidance. This tilts the balance such that, when an odor is presented, approach MBONs respond at a higher rate than avoid MBONs, driving approach behavior (Fig. 7A*↔*B, initial training). Avoidance behavior operates in the opposite way by weakening CS-responsive KCs to approach MBON synapses. With this model, we asked, given a test trial, what are the contributions of reliable and unreliable KCs towards overall discrimination?

**Figure 7:**
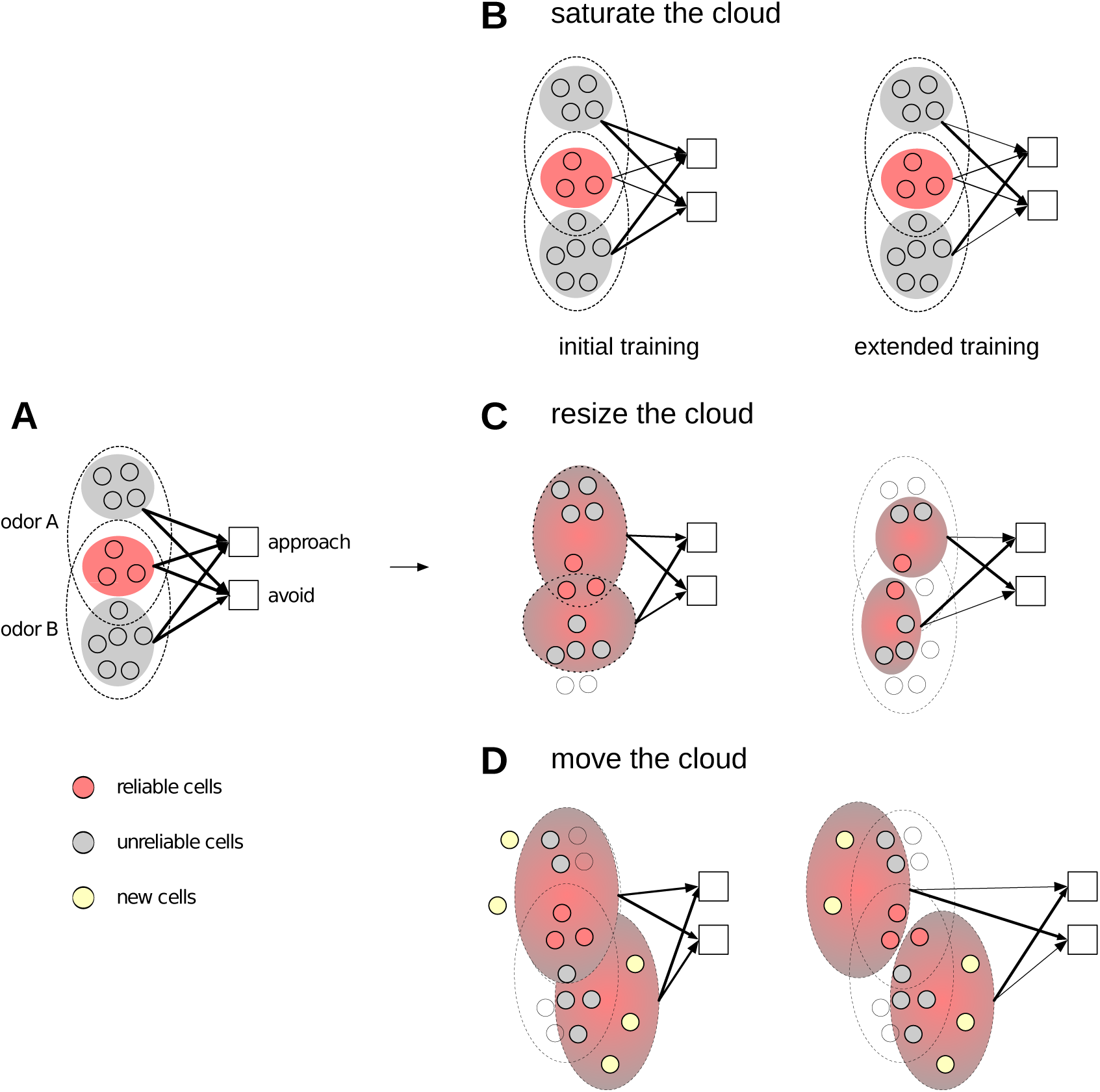
Three possible models of fine-discrimination. (A) The stimulus representations in the untrained state. For each odor, the dotted ovals enclose the cells that respond to the odor. These cells comprise unreliable cells (grey) and reliable cells (pink). The unreliable cells are mostly non-overlapping for the two odors, whereas the reliable cells overlap exactly. The thickness of the arrows denotes the total strength from the three sets of cells to the two downstream neurons that drive approach or avoidance behavior. Initially, all arrows are of the same strength. (B) Saturating the cloud. With initial training (left), the strength of downstream synapses of reliable cells saturates (i.e., weakens to near zero), but synapses of unreliable cells barely change their strength. Consequently, as reliable cell overlap is high between the two odors, approach/avoid neurons receive equal input, and the animal is unable to discriminate. With extended training (right), the downstream synapses of unreliable cells also saturate, driving the two stimuli apart, as depicted by the thicker arrows. Consider the top odors connections (arrows) strengths to approach and avoid MBONs. The connection to approach are much weaker than avoid, and therefore odor A and B are distinguishable. (C) Resize the cloud. The stimulus representation shrinks to only include cells that are non-overlapping between the two stimuli (e.g., shared reliable cells are turned off). While some resizing may occur with normal training, with extended training, the two stimuli are further driven apart, enabling the animal to discriminate the two odors. (D) Move the cloud. Some mechanism (most likely driven by unreliable cells, the cells that are most unlike between similar odors) rearranges the stimulus ensemble response to include new cells or converts unreliable to reliable cells that do not overlap, and thus improve discrimination.

We illustrate the answer by considering the case where odor *A* is trained with a reward and odor *B* is trained with punishment, and pick an example KC *x*, which responds to *A* in 2 trials (out of 6 total trials) and to *B* in 4 trials. Thus, *x*’s response probability is 1/3 for *A* and 2/3 for *B*.

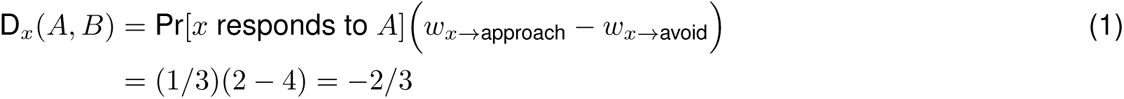

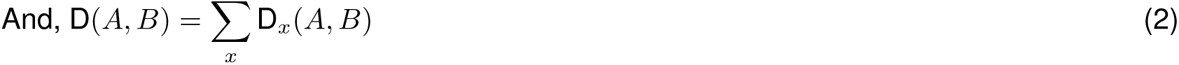

There are 3 equations that effectively describe the contribution of *x* and all KCs towards discrimination, when *A* is presented in a test trial. Intuitively, the contribution of *x* depends on how its response differs between both odors. Equation 1 describes the contributions (D*_x_*) of *x* towards approach, and the total discriminability of odor *A* from odor *B* is simply the sum of D*_x_* over all *x* (eq. 2). The first factor (1*/*3) in eq. 1 represents the probability of activating cell *x* when *A* is presented. The second factor represents the difference in the synaptic weights (*w_x_*’s) of the connections from *x* to approach and avoid. The *w_x_*’s are determined by equation 3.

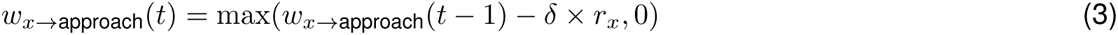

Equation 3 outlines how the weights *w_x→_*_approach_ and *w_x→_*_avoid_ (initially high and equal) change with each training trial. Consider exposure to A and reward: *w_x→_*_avoid_ decreases in proportion to the size of *x*’s response (*r_x_*) and learning rate *δ*. The same rules apply to exposure to B and punishment and *w_x→_*_approach_. The *t* variable represents trial number or time, and shows that *w_x_* will be updated differently depending on whether it is a reliable or unreliable KC synapse. The depression in synaptic weights is faster for reliable cells as they are active more often (83% of the time, Fig. 1) with larger responses. Moreover, as synapses cannot decrease beyond 0 (0 term in eq. 3, saturation of synapses), reliable KC synapses will approach 0 quickly, i.e., after very few trials. By contrast, unreliable cell synapses will require many trials to reach 0–weight as they are active less often (25% of the time, Fig. 1) with smaller responses.

The *w_x_*’s (of reliable and unreliable cells) also depend on the amount of training they undergo. As per Chapuis & Wilson (2012), normal training is the shorter training period required for discriminating dissimilar odors (e.g., 6 trials for flies (Tully & Quinn, 1985; Campbell et al., 2013)), and extended training is the longer training required to distinguish similar odors (e.g., 100 trials). Revisiting equation 1, let us assess the effect of a normal training regimen. D*_x_* depends on whether *x* is a reliable, unreliable, or silent for odors A and B. Table 1 lists the possibilities. D*_x_* is large when *x* is reliable for one odor but not the other (lines 2–4). With extended training, if *x* is responsive to both odors (whether as R or U), D*_x_* will be close to 0 as reliable and unreliable synapses will have saturated. Thus, D*_x_* is large only when *x* is silent for B. Since there are many more unreliable cells and they are unlikely to overlap, the overall D increases with extended training.

**Table 1:** Possible contributions by cell *x* towards discrimination. Columns 1 and 2 show *x*’s response state (R–reliable, U– unreliable, or 0–silent) to odors A and B. Columns 3 & 4 show the discrimination contributions for normal – N(D*_x_*) – and extended – E(D*_x_*) – training. As we are interested in contributions that distinguish A from B, *x*’s silent for A are ignored. For normal training, rows 2–4 generate high contributions, while rows 1 & 5 have low contributions. With extended training, only cells silent for B generate high contributions.

| Row | A | B | N($D_x$) | E($D_x$) |
| --- | --- | --- | --- | --- |
| 1 | R | R | $R - R \sim 0$ | 0 |
| 2 | R | U | $R - U \gg 0$ | 0 |
| 3 | R/U | 0 | $R \gg 0$ or $U \sim 0$ | $\gg 0$ |
| 4 | U | R | $U - R \ll 0$ | 0 |
| 5 | U | U | $U - U \sim 0$ | 0 |

The results of applying MB/PCx responses from Figure 1 to equations 1–3 are shown in Figure 6. In it, dissimilar odorpairs are those with less than 0.15 correlation in their KC responses (Fig. 4), and similar ones are those above 0.5. Three features become apparent for both species. First, with (N)ormal training, the contribution (D*_x_*) of unreliable cells (*x*) is about the same for all odor–pairs (flies: 1.2 vs. 1.2; mice: 2.7 vs. 1.8), while reliable cell contributions decreases from dissimilar to similar (flies: 4.1 vs. 1.1; mice: 3.6 vs. 1.7), because reliable cells for similar odors are highly likely to overlap. This result reflects observations of similar odor discrimination being a hard task (Campbell et al., 2013; Lin et al., 2014; Chapuis & Wilson, 2012). Second, as reliable cell synpases are already saturated with (N)ormal training, their contributions do not increase for (E)xtended training (flies: 4.6 vs. 4.1 for dissimilar odors and 1.6 to 1.1 for similar odors; mice: 3.6 vs. 3 for dissimilar and 1.7 vs. 0.4 for similar). Note that some reliable cells even have reduced contributions – as shown in row 2 of Table 1 – with extended training, when their response to odor B is unreliable. Third, unreliable cell contributions increase with extended training (flies: 1.2 vs. 4.2 for dissimilar and 1.2 to 4.1 for similar; mice: 2.9 vs. 10 for dissimilar and 1.7 vs. 1.7 vs. 7.6 for similar).

Thus, with normal training, the discrimination measure for the whole population is reduced from dissimilar to similar odor pairs because of the reduced contributions of reliable cells owing to the correlation in their responses to similar odors. With extended training, however, the discrimination measure for the population goes up again as the contribution of unreliable cells (which are decorrelated irrespective of odor similarity) goes up.

These analyses show that reliable and unreliable cells may encode different kinds of information, which are both useful towards identifying odors. As an illustration, let’s say the target odor (*A*) belongs to the citrus family of fruits, and the comparison odors (*B*’s) all belong to the berry family. In this case, reliable cells provide distinguishing information, because, for dissimilar odors, reliable KCs are decorrelated. On the other hand, if the comparison odors also belong to the citrus family, then reliable KCs provide less distinguishing information, since reliable KCs are highly correlated for similar odors. Unreliable KCs, however, can provide distinguishing information, as they are decorrelated for similar odors (Fig. 4C). Thus, we propose that reliable and unreliable cells may provide different levels of odor identity information, with reliable KCs providing first-order identity information, and unreliable KCs providing second-order identity information, which becomes important as odors become similar, or which may encode finer, more subtle features of the odor.

## Discussion

### Summary of results

In this manuscript, we showed that higher sensory regions responsible for odor discrimination and learning, in both flies and mammals, encode odors using a stochastic code. The stochastic code comprises cells that fall along a continuum based on the cell’s reliability (i.e., the fraction of odor trials for which the cell responds). Reliable cells respond in most odor trials with a larger responses. By contrast, unreliable cells respond in fewer trials with smaller responses. With a firing rate model of the fly olfactory circuit, we show that the major driver of the stochastic code is synaptic noise in the winner-take-all circuit between the APL neuron and KCs, and is unlikely to be a product of sensory noise alone. Finally, using the fly olfactory association paradigm, we show that such a stochastic code can enhance discrimination ability.

Building on this experimental observation, below we: 1) discuss insights into neural circuit mechanisms that give rise to stochastic codes; 2) hypothesize benefits for stochastic codes and propose models for how reliable and unreliable cells work together to facilitate odor discrimination; 3) hypothesize possible mechanisms in flies and mice; 4) revisit the role of sparse coding for fine-grained discrimination; and 5) describe other neural circuits and brain regions where benefits of trial-to-trial variability have been appreciated.

### Neural circuit mechanisms for generating stochastic codes

There are two inputs that shape the activity of an odor-coding cell in MB: inputs from PNs and from APL. First, each Kenyon cell (KC) samples from approximately 6 of the 50 PN types, and this input plays a role in whether the cell is reliable or unreliable. For example, a KC is very likely to be reliable for an odor if most of the 6 PNs it samples from are highly active (large response) for the odor. This would allow the Kenyon cell to survive any reasonable amount of noise or variance in the winner-take-all threshold. On the other hand, a KC is more likely to be unreliable if only half of the PNs it samples from are highly active. Combinatorially, there are many distinct odors for which only three or fewer out of six PNs are highly active. Similarly, if a specific PN*→*KC synapse is strong, then perhaps only that one PN need be strongly active for an odor for the KC to be reliable, but there are likely many more odors where that PN is mildly active. Thus, for any odor, there are many more unreliable cells than reliable cells.

Considering the stochastic component, we experimented with three possible causes of variability with our model. Two of the causes were based on sensory noise arising in the first or second stages of the circuit (i.e., noise in PN firing rates or in PN*→*KC synaptic transmission). For both, it was implausible to generate the observed stochasticity without physiologically excessive noise levels. Previous studies support this view (Srinivasan et al., 2018), showing that in the mouse PCx circuit, each neuron receives synapses or inputs from many OB neurons, and noise reductions in one input are likely to be offset by noise additions in other inputs. The fly has fewer inputs into each KC in comparison, but even here, averaging is likely to filter some noise. In addition, connectome analyses of MB have found compensatory variability in the fly circuit; e.g., weights of excitatory PN*→*KC connections are inversely correlated with the number of PNs each KC samples from (Abdelrahman et al., 2021), effectively canceling out variability. Thus, for moderate noise regimes, odor signals are robustly transmitted to the third stage of the circuit making it unlikely that the observed coding variability originates from the first two stages alone.

The WTA circuit, on the other hand, does not average or cancel out noise. Each Kenyon cell gets direct inhibition from APL, and thus, is sensitive to variation in the APL*→*KC synapse weight. Computationally, this implies that in each odor trial, each KC *x* receives a slightly different amount of inhibition, which alters the relation between its firing rate, *r_x_*, and its threshold for activation, *τ_x_*. For example, KC *x* could receive inhibition sampled from a (0-mean) distribution with per-KC variance *σ_x_*per trial. If *r_x_ ≫ τ_x_*, then KC *x* survives inhibition regardless of the variance, and these are the reliable KCs. KCs hovering around *τ_x_*will only fire during some trials, and these are unreliable KCs.

The mouse PCx, like the fly MB circuit, contains a WTA circuit between PCx principal and inhibitory cells (albeit in a different form (Bekkers & Suzuki, 2013; Franks et al., 2011; Bolding & Franks, 2018)), providing multiple synapses where variability can arise. Variability may also arise through other sub-circuits in PCx. One of them is the excitatory recurrent circuit between PCx principal cells (Franks et al., 2011), which accounts for nearly half of excitatory PCx principal cell inputs (Srinivasan et al., 2018; Sarma et al., 2010). Additionally, olfactory bulb input into layer 2 and 3 cells is modulated by a feedforward inhibitory circuit between mitral cells in OB and principal cells in PCx (Suzuki & Bekkers, 2010; Bekkers & Suzuki, 2013). Finally, PCx cell activity is likely influenced by an OB*→*PCx*→*OB feedback circuit that has the characteristics of a WTA circuit, in negatively suppressing its own activity. A portion of PCx principal cells (which are activated by OB mitral cells) excite granule cells in the bulb that then inhibit mitral cells (Otazu et al., 2015; Chu et al., 2016; Yamada et al., 2017), and, could, in turn, change OB input to PCx, and PCx activity.

Thus, processes that convey sensory information from the periphery collaborate with inherent circuit mechanisms within MB and PCx to encode odor responses using a stochastic code. The contribution of the winner-take-all circuit, as opposed to input noise, highlights an intriguing possibility of neural circuit design. All circuits might contain mechanisms to filter noise from upstream circuits, while also containing mechanisms that generate inherent noise that aid in better exploration of the environment, or as we suggest next, better discrimination.

Finally, our study is compatible with work showing that odor identity is also encoded by a temporal or primacy code (Wilson et al., 2017; Kepple et al., 2019), in which identity is derived from the combination of neurons that respond the earliest (Stern et al., 2018). Reliable cells, which respond more frequently with larger responses (spikes/s), would have an earlier response than unreliable cells. Recent findings (Bolding & Franks, 2017, 2018) showing that fine-discrimination requires more time, with the PCx WTA circuit facilitating this discrimination, support this view and are consistent with our findings.

### Benefits of stochastic coding, and possible models for fine-grained discrimination

Perceptual learning is the phenomenon where an animal is unable to distinguish two similar stimuli but after repeated (passive) exposure or (active) training sessions, acquires the ability to do so (Chapuis & Wilson, 2012; Green et al., 2018). Responding cells for two dissimilar odors will be mostly non-overlapping, which means that reliable cells are by themselves likely to be sufficient for distinguishing dissimilar odors. However, given two similar odors, we showed that there is stronger correlation in the activities of reliable cells than the activities of unreliable cells (Figs. 2,4). Thus, unreliable cells may encode important information (e.g., higher order statistics or more subtle features of the odor) that can, with training, help discriminate the odors. Our model (Fig. 6) shows that it is harder to discriminate similar odors with normal training, and extended training enables better discrimination by recognizing the information encoded within unreliable cells. While the question of how much distinguishing information MB or PCx needs to discriminate two odors remains unknown — e.g., is a single non-overlapping cell sufficient? — it is generally accepted that the more non-overlapping the representations of two odors, the easier it is to discriminate (Cayco-Gajic & Silver, 2019).

Below, we describe three possible models for fine discrimination (Fig. 7), and for the purpose of illustration, we use the fly olfactory learning system (Modi et al., 2020). These models transition from thinking of an odor as a single point in highdimensional space to thinking of an odor as a cloud in high-dimensional space. Each point in the cloud corresponds to the representation assigned to an odor in a single trial. In addition, these models require extended training, with many more training trials than is typically performed, since unreliable cells are active in *≤* 50% of trials.

To begin, Figure 7A depicts two similar odors, A and B. The responsive neurons for A and B fall into three clouds. The pink cloud contains the reliable cells, which are highly overlapping since the two odors are very similar. The unreliable cells are less overlapping and are depicted by two gray clouds for odors A and B. Since the system has not undergone training, the strength of the connections that drive downstream approach and avoid MBONs are equally matched. Thus, the fly neither approaches nor avoids the odor upon exposure.

*1. Saturate the cloud (Fig. 7B)*. The first model performs perceptual learning by saturating synapse strengths between unreliable KCs and MBONs. The results of Figure 6 show this model in action. First, with initial training — e.g., 12 CS-US trials or a similar amount as in Campbell et al. (2013), which could be considered as a norm for most studies — the strength of synapses from reliable cells (for the two odors being compared) to MBONs are depressed. Second, the strength of synapses from an odor’s unreliable cells to the MBON are depressed, however, not as much as reliable cell synapses, since unreliable cells are activated in fewer trials. For similar odors, the reliable cells are a “wash”, providing little information for distinguishing odors, whereas unreliable cells have begun to separate the odors, though they have not responded in enough trials to make a robust behavioral difference. With extended training, the effect of learning on unreliable cell synapses increases to make a difference and aid discrimination.
*2. Resize the cloud (Fig. 7C).* In the second model, the cells encoding the two odors change with training, such that the cloud for odor A is resized to exclude cells that also respond to odor B. The cloud can change size by converting unreliable cells to reliable cells or by turning off reliable cells (since they overlap the most between two similar odors).
*3. Move the cloud (Fig. 7D).* The third model moves the clouds apart via the addition of new cells (colored in yellow) that were not previously responsive to the odor. In addition, unreliable cells may become more reliable with extended training.

Finally, there remains an important unresolved problem: both saturating the cloud and moving the cloud could potentially cause unintended interference with other odors. For example, when saturating the cloud, unreliable cells for odor A now strongly drive approach behavior; however, these same unreliable cells could be reliable cells for odor C, and if odor C was paired with punishment, there could be a problem. Similarly, when moving the cloud, the new cells recruited to be part of odor A’s cloud could previously have been part of another odor’s cloud. With all models, although training improves the discriminability of a particular odor–pair, it will affect the discrimination of other odors and the total number of odors that can be distinguished (odor capacity). Thus, in future experimental evaluations of these models, it is important to consider how well stable odor perception is maintained and how coding capacity is affected.

### Learning and discrimination mechanisms in piriform cortex and mushroom body

Although, the models sketched here need to be grounded in precise circuit mechanisms, there is evidence for cloud saturation in flies, and evidence in favor of cloud saturation and cloud moving in mice. In flies, previous experiments that used optogenetic activation of DANs as reinforcement (Hige et al., 2015) showed that KC representations and amplitudes remain stable with associative training, while MBON activity decreased. While there may be other effects during extended training (e.g., non-associative effects based on anatomical or neuromodulatory feedback from MB to the antenna), these results support the “saturate the cloud” model in flies (Fig. 7B).

In vertebrates, there is evidence that cell responses remain stable with learning in mouse (Wang et al., 2020), while other studies in rat PCx (Shakhawat et al., 2014), mouse PCx (Schoonover et al., 2021), and fish Dorsal Pallium (Jacobson et al., 2018) show that they change over time (moving the cloud model). Intriguingly, Schoonover et al. (2021) showed that *∼* 2.5% of PCx cells remained stable. Based on our findings of 3.3% reliable cells, it is likely that these are reliable cells, and might explain the contradictory results observed; while reliable cells remain stable, unreliable cells change over time. Even though unreliable cells change, their effect on discrimination would still be compatible with our model (Fig. 6) as long as the drift for odor-pairs is decorrelated. The notion that PCx cells play a part discrimination comes from an important study examining the discrimination of artificial PCx odors (Choi et al., 2011) — generated by using light on PCx neurons. The ability of PCx ensemble activity (sans sensory activity) to generate discriminatory behavior suggests that learning and discrimination mechanisms are likely in PCx or downstream of it. If they are downstream, in which region could the plasticity mechanisms operate? Several labs have shown that OB responses decorrelate over time, and that PCx seems to be required in the form of PCx *→* granule cell inhibition of mitral cells (Otazu et al., 2015; Yamada et al., 2017; Boyd et al., 2012; Kato et al., 2012), with some evidence of plasticity in OB (Sailor et al., 2016). Notably, OB decorrelation occurs even without reinforcement (which is unlike the fly mechanism). Additionally, if OB encoding changes over time, it might also lead to changes in PCx coding of odors favoring the moving the cloud model. Other candidate downstream areas that might do the work of the KC-MBON-DAN circuit include the Posterior PCx or parts of the prefrontal cortex or even the hippocampal formation (Wang et al., 2020; Calu et al., 2007; Meissner-Bernard et al., 2019; Woods et al., 2020), but further studies are required.

Thus, learning and discrimination mechanisms in the olfactory circuit could be a hybrid scheme that includes saturation of downstream synapses and changes in odor coding ensembles. We are not aware of cloud resizing in the olfactory system; however, there is evidence of stimulus ensemble resizing in the context of songbird and motor learning, which we discuss below.

### Challenging the role of sparse coding

It has long been argued that increased sparsity in odor representations leads to increased pattern separation between two odors (Cayco-Gajic & Silver, 2019). While sparse coding may, indeed, contribute to discrimination of dissimilar odors, sparse coding alone may not be sufficient for fine-grained discrimination of two very similar odors. The highest firing cells (i.e., those that would remain firing if odor representations were made sparser by increasing inhibition) would be reliable cells, which have more overlap, not less, between similar odors (Figs. 5,S8). An intriguing alternative possibility, for flies, is that the inability to discriminate between similar odors arises from the insufficient effect of unreliable cells, as they do not experience enough trials under standard training protocols.

Why, then, is there a need for sparse coding? There are two potential uses for it. First, sparse coding is integral to discrimination, not in separating two odors, but in separating many odors; i.e., for enhancing the coding capacity of the system. When odor codes are sparse, they use fewer cells, and thus when new odors are introduced, they are less likely to interfere with stored odors. For example, if an odor is encoded by 5% of Kenyon cells, and we conservatively assume that new odors can be learned only if they are completely non-overlapping with pre-existing odors, then 20 odors can be learned. If the odor code were less sparse, at say 10%, the capacity of the circuit would be lower at 10. While actual discrimination may be more forgiving with respect to odor overlap, the notion that coding capacity is inversely related to sparse code size still applies. Sparse coding may be important for optimizing the coding capacity of the circuit but our results suggest that it may not improve discrimination between similar stimulus pairs. Further, there is a limit to how sparse a code can be. Odor recognition using very sparse codes (e.g., if only one or two neurons were to encode odors) is likely less robust to sensory noise and other environmental nuisances. Second, while sparse coding might not aid discrimination of similar odors, it reduces the computational power needed for dissimilar odors, which might have an evolutionary benefit. Thus the olfactory system may be geared towards quick discrimination using sparse coding (reliable cells), and towards longer discrimination of similar odors by leveraging unreliable cells.

### Generality to other neural circuits and brain regions

Trial-to-trail variability in neural responses is common in myriad sensory regions and species, and other studies have shown at least three ways in which variability can improve an animal’s fitness.

First, variability expands the global search space. A popular illustration of this phenomenon is in songbirds (Tumer & Brainard, 2007; Woolley & Kao, 2015). Young zebrafinches learn their courtship song from older adults in a process called directed singing. They also engage in solo practice sessions called undirected singing, where they vary the song. The final song used in courting females is a mix of the stereotyped song that they learned and the variable song that they practiced. Related theoretical studies, especially in computer science, have shown that a crucial part of reinforcement learning algorithms is a random exploration component (Sutton & Barto, 1998), which helps guide the search away from local minima. The use of variability in song learning is an instantiation of this principle. A similar strategy is also observed in motor systems. The amount of learning required is directly proportional to the amount of variability at the start, and as learning or training continues, the amount of variability is reduced (Dhawale et al., 2017). Essentially, the subject explores various possibilities at the start, and then exploits the best solutions found. Here again, variability increases the spread of possible actions and thus the likelihood of hitting an optimal solution. Thus, for olfaction, it is intriguing to hypothesize that variability in odor coding may vary over time, perhaps tuned by neuromodulators, until some “optimal” encoding is found. Notably, this notion of improving the search for a global solution finds some support in theoretical studies of neuronal networks. Neuronal networks formed in the presence of noise are more robust and explore more states, enabling them to better adapt to dynamically changing environments (Krogh & Hertz, 1992; Kirkpatrick et al., 1983).

A second use of variability is the phenomenon of stochastic resonance (SR) found in various species and systems (Longtin et al., 1991; Cordo et al., 1996; Levin & Miller, 1996; Douglass et al., 1993). The cricket cercal sensory neurons (Levin & Miller, 1996), and crayfish mechanoreceptors (Douglass et al., 1993) provide a good illustration of how SR is useful. In both, the ability of animals to sense other animals or objects in their vicinity is dependent on the threshold activation of their sensory receptors: the lower the sensitivity threshold, the farther the range for sensing a signal. Extremely small thresholds for sensing a signal, however, are problematic, as animals might then confuse noise and signal. In an interesting twist, noise presents a solution. If high thresholds are augmented by random noise events, the effectively lowered thresholds can allow the animal to sense better, while alleviating the signal being mistaken for noise issue of using low thresholds. This kind of detection allows the system to carry more information, as the output of the detector neuron reflects all possible levels of activity, not just signals above the threshold (Shannon, 1948). More studies are needed to understand the noise regime that can modulate receptor activation while balancing the tradeoff between easy detection and keeping signal–to–noise ratio low.

The WTA mechanism resembles these SR mechanisms: KC responses are thresholded (otherwise all KCs would respond), and there is evidence that noise affects the activation threshold on an infrequent basis (unreliable cells). Noise that effectively lowers the threshold would improve detection of the odor. The improved detection, in turn, favors better discrimination, because the noise-induced (unreliable) KCs are unlikely to overlap with other odors. The WTA mechanism, thus, presents evidence for SR in a non-peripheral circuit, unlike earlier instances of SR.

In this study, we suggest a third, hitherto, unexplored benefit that might accrue from trial-to-trial variability: the ability to improve learning and discrimination, specifically for fine grained discrimination.

Overall, our observation of broad conservation between the two circuits leads us to believe that a stochastic code arising through the action of a WTA mechanism within a distributed circuit might be more universal. Other distributed circuits such as the hippocampus, prefrontal cortex, and the cerebellum have prominent WTA circuits (Cayco-Gajic & Silver, 2019), with observed instances of trial variability. Circuit specific differences would explain differences in responses and also help in elucidating the adaptation of stochasticity for circuit-specific needs.

## Supplement

### Figures

**Figure S1:**
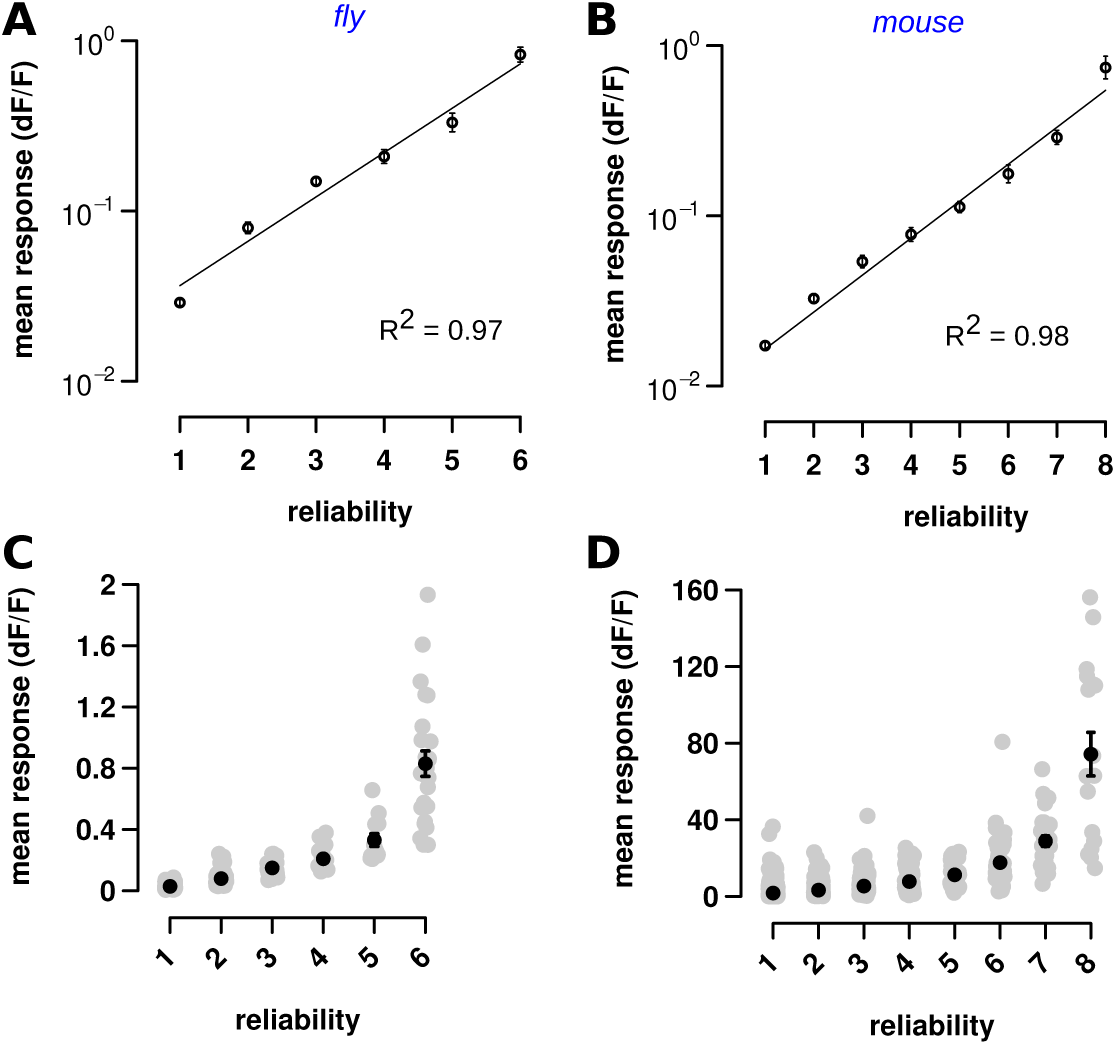
Odor response sizes in the fly MB and mouse PCx cells increases with reliability. (A-D) Mean response size of each cell plotted against its reliability level (on the x-axis) for fly (A,C) and mouse (B,D). (A,B) Shows the fits for the curves in Figures 1C,F. In both cases the exponential equation y = aebx has been converted to the form log(y) = log(a) + bx in order to do a least squares linear regression fit, with the fit r shown in the plots. Error bars are mean ± SEM, and have been converted to a log scale taking care that the log error is properly propagated. (C,D) The two figures are similar to Figs. 1C,F, except here all the points are plotted instead of just the means. Each grey circle denotes a cell’s response to an odor on the x-axis, and represents the mean response size of that cell on the y-axis. The reliability number on the x-axis denotes the number of trials (reliability) for which that cell had significant responses. Error bars are mean ± SEM.

**Figure S2:**
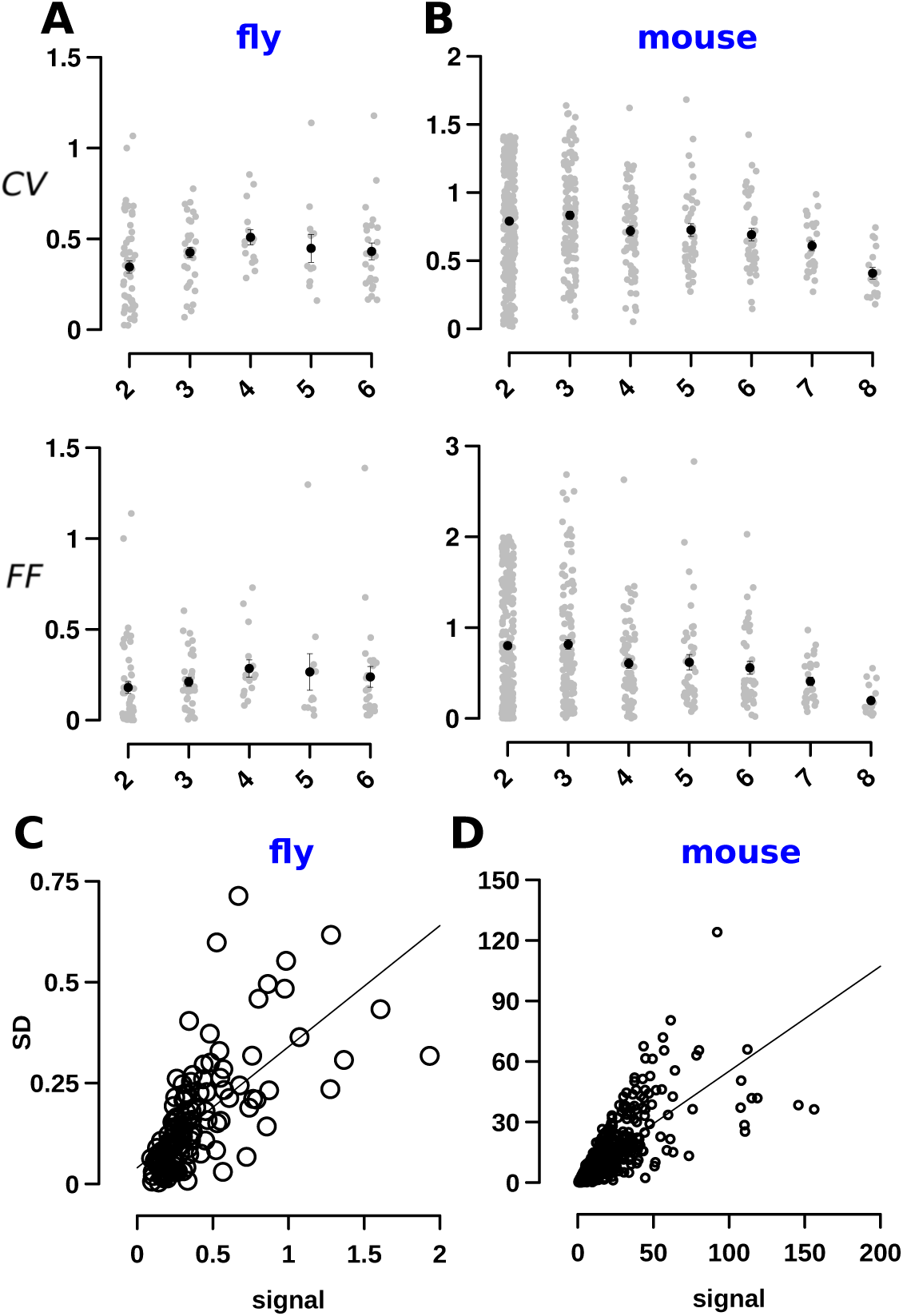
Responses in Mouse PCx cells are more variable. (A) Coefficient of Variation (CV) and fano factor (FF) of cell responses in flies. (A, top) shows CV (on the y-axis) with cells split according to their reliability (on the x-axis). The plot does not include cells with a reliability of 1. There is no significant difference in CV across different reliabilities. Error bars in this and all the plots are SEM. (A, bottom) FF for the fly showing how FF differs depending on cell reliabilities. FF for the different cell reliabilities are not significantly different. (B, top) CV for the mouse plotted for different cell reliabilities. CV is greater for the mouse compared to the fly, and decreases with lower cell reliabilities. (B, bottom) FF for the mouse plotted against cell reliabilities shows that FF is higher in the mouse than fly, and FF decreases with cell reliabilities. (C,D) Plot showing how the standard deviation (SD) varies with response magnitude. In both, flies, and mice, SD increases with increase in signal, and are fit for lines with slopes of 0.52 and 0.5. The units are a normalized readout of Ca2+ based flourescent activity.

**Figure S3:**
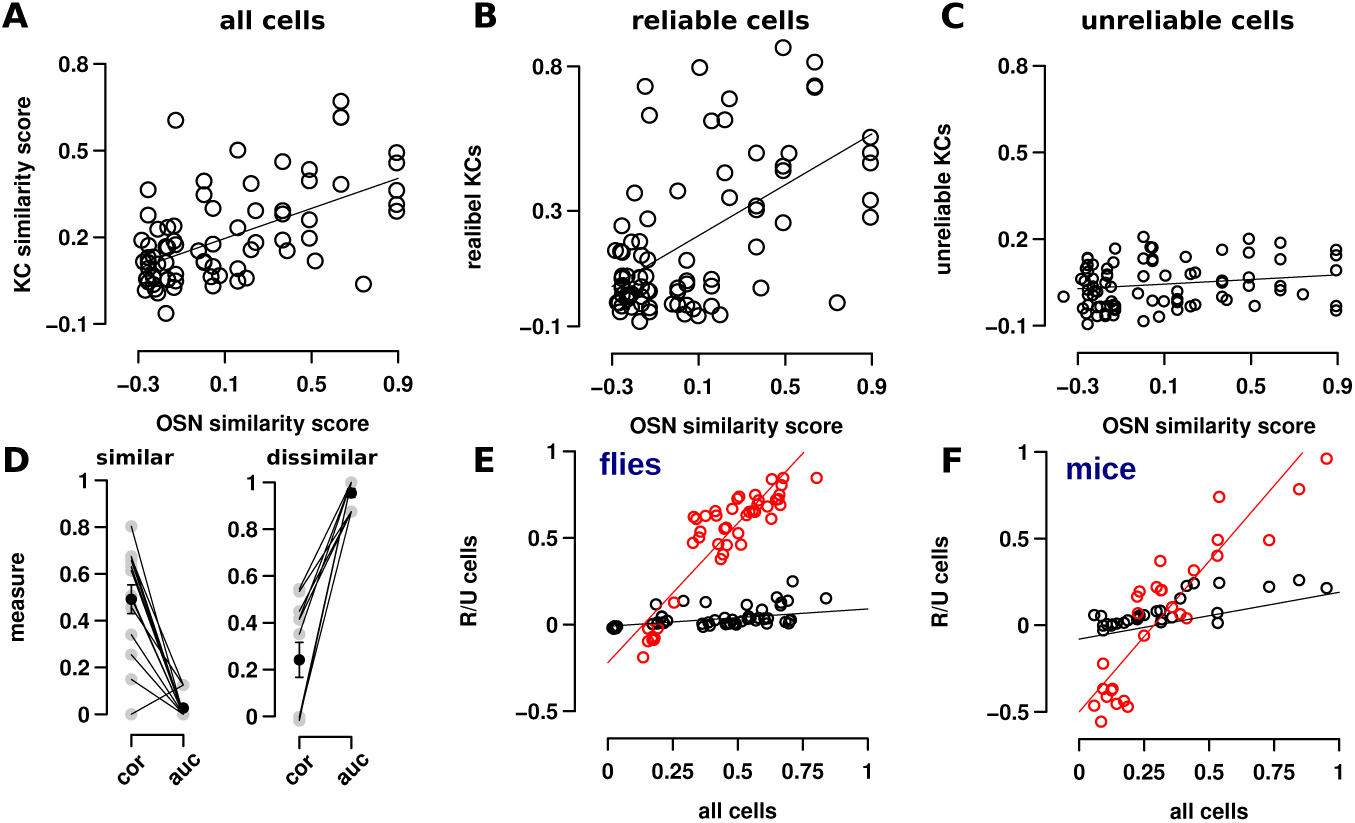
Reliable cells better preserve odor–similarity. (A) The similarity between all odor pairs in the fly’s nose (Hallem et al., 2006) (x-axis) and MB (y-axis) fits the line fit y = 0.17 + 0.44x (R2=0.57). Similarity is measured as the correlation between odor-pairs, and each circle denotes one odor-pair. (B) The similarity between odors pair responses in the fly’s nose and MB judged by reliable cells alone is a fit for the line fit y = 0.16 + 0.32x (r=0.5). On the other hand, the correlation between fly’s nose and MB is low when judged by unreliable cells alone in (C). The line fit has a slope close to 0 (y = 0.04 + 0.04x, r=0.2), though showing a slight trend, wherein odors that are dissimilar (to the left of the x-axis) have less correlation than odors that are similar (to the right of the x-axis). This figure shows similarities for all odor pairs in the fly’s antenna and MB, unlike the main figure (Fig. 4) which is restricted to those odor pairs for which correlations in the antenna and MB are within 0.2 of each other. (D) AUC measures the degree to which odors are distinguishable. Comparison of the AUC measure with correlations. In the left panel, are all odor-pairs that have an AUC measure below 0.25 (mean: 0.04) have a corresponding odor similarity of 0.5 (mean) as measured by Pearson’s correlation. In the right panel, odor-pairs that have an AUC measure above 0.75 (mean: 0.93) have a corresponding odor similarity of 0.24. (E,F) Similarity of odor-pairs based on all cell populations versus just the population of (R)eliable (in red) or (U)nrelibale (black) cells for both flies and mice. In both species, while Reliable cell similarity is proportional to odor–pair similarity of all cells, the Unreliable cell similarity remains low for all odor–pairs.

**Figure S4:**
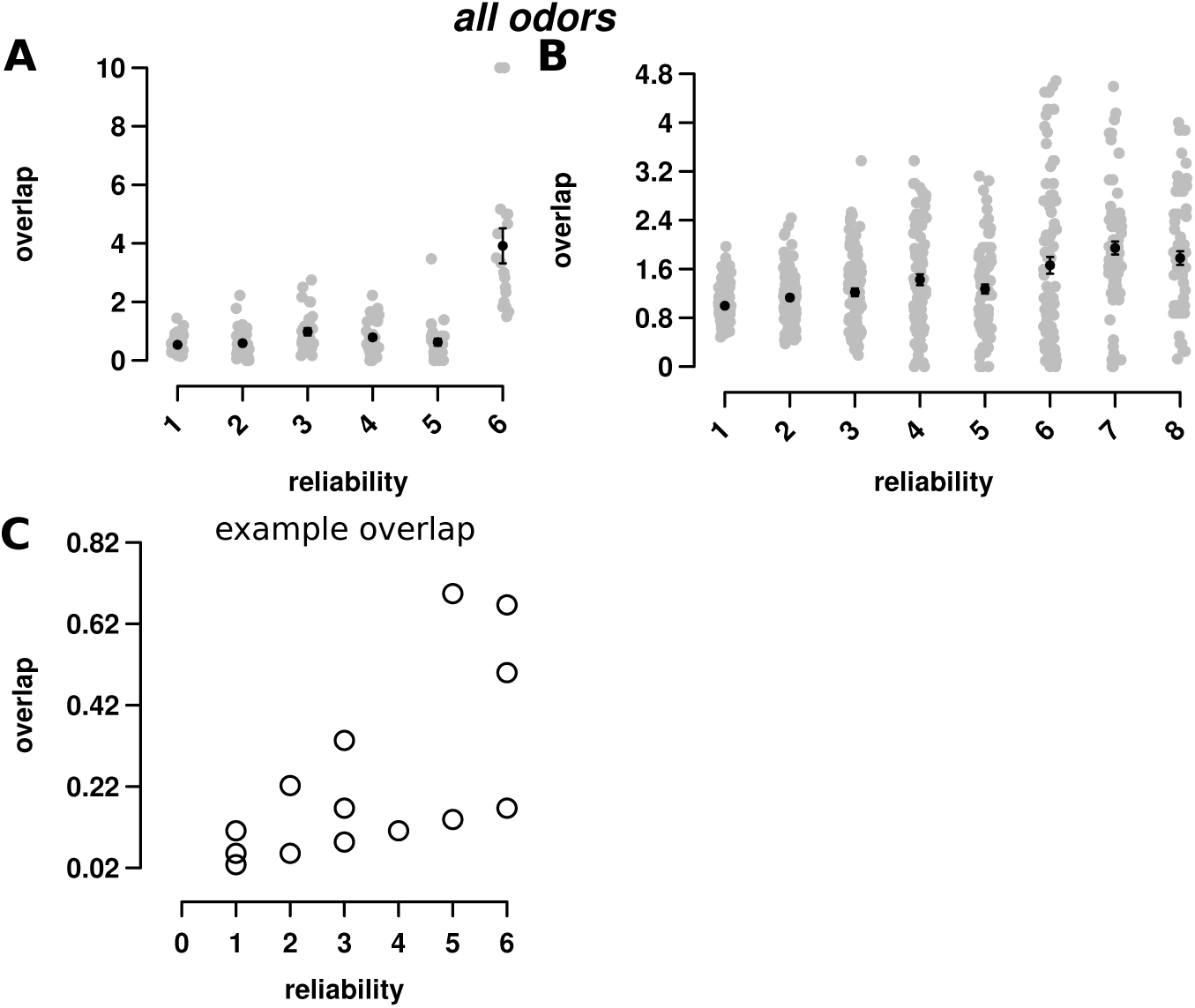
Unreliable cells are likelier to respond differently between similar odors. (A,B) Reliability versus probability of overlap plots for flies (A) and mice (B). For each point or reliability on the x-axis, we plot the corresponding overlap for all odor-pairs while only considering cells with that level of reliability. In both plots, unreliable and reliable cells have similar amounts of overlap, although cumulatively, reliable cells have slightly higher average probability of overlap. (C) An example of using an overlap measure to assess the similarity of odors. As the reliability on the x-axis increases a cell’s overlap with the other odor increases. Each circle denotes a single cell and it’s overlap measure calculated as the probability of response to odor 1 * probability of response for odor 2.

**Figure S5:**
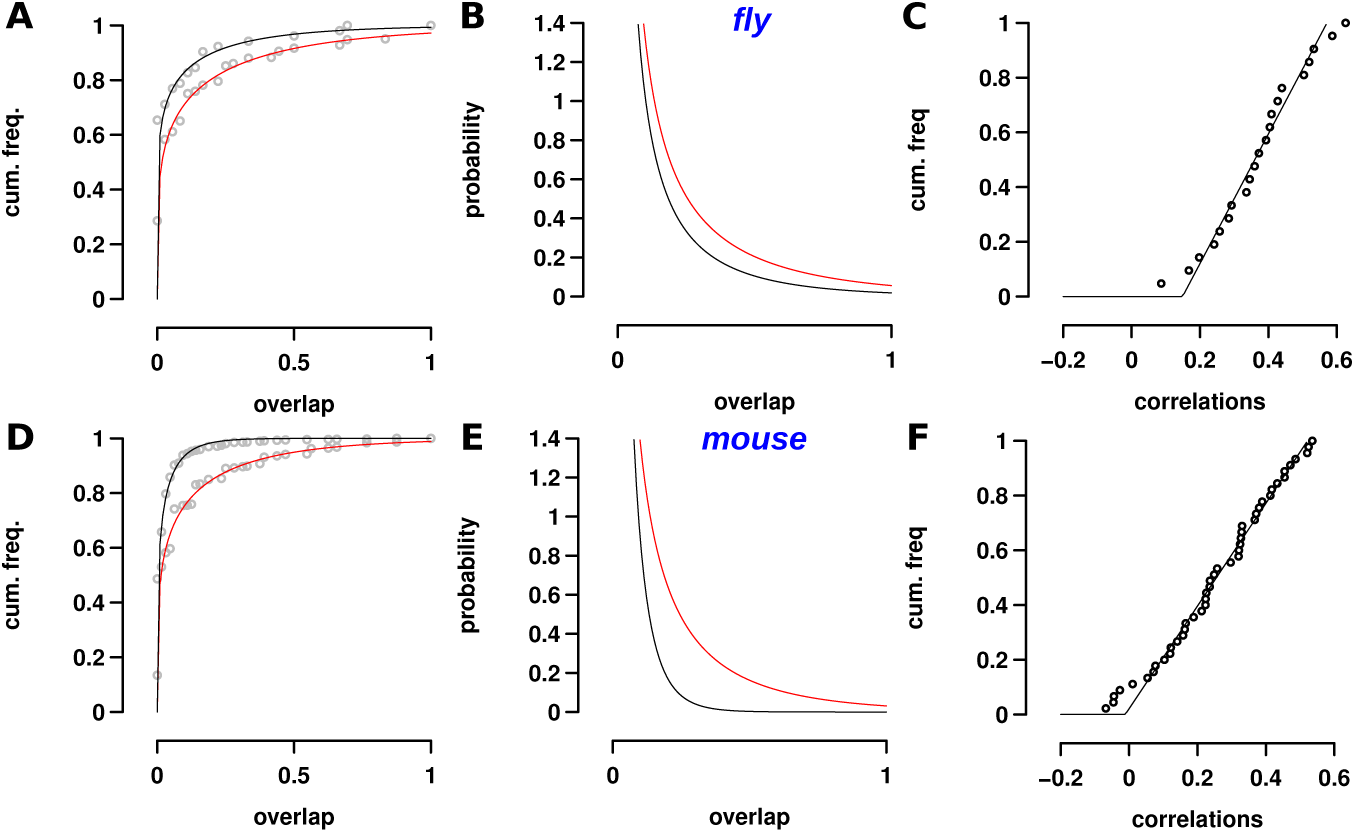
The probability of overlap cell increases from similar to dissimilar odors. (A-C) Comparison of overlap statistics between similar and dissimilar odors for flies. (A) Shows the cumulative frequency for overlaps between cells for dissimilar (top curve) and similar (bottom curve) odor-pairs. Similar odors were odor-pairs with correlation > 0.5, and dissimilar odor-pairs had correlation < 0.15. The plot shows that for low overlap measures – extreme left – both types of odor-pairs have similar probabilities. But, the probability of higher overlap increases for similar odors. To illustrate the point, we also show the PDF in (B), with similar odor PDFs shown in red. (C) The cumulative distribution of correlations between all odor-pairs shows that correlations are uniformly distributed in the interval (0.14,0.57). (D-F) Comparison of overlap statistics between similar (bottom curve in D and red curve in E) and dissimilar odors for mice. (D,E) Comparing the overlap distribution for dissimilar and similar odors shows that while the probability of cells having a low overlap is the same, the probability of cells having a high overlap is much higher for similar odors. (F) The similarity between odors is evenly distributed in the dataset that we analyzed. The plot shows the cumulative probability distribution is a good fit for a uniform distribution on the interval (–0.01,0.52).

**Figure S6:**
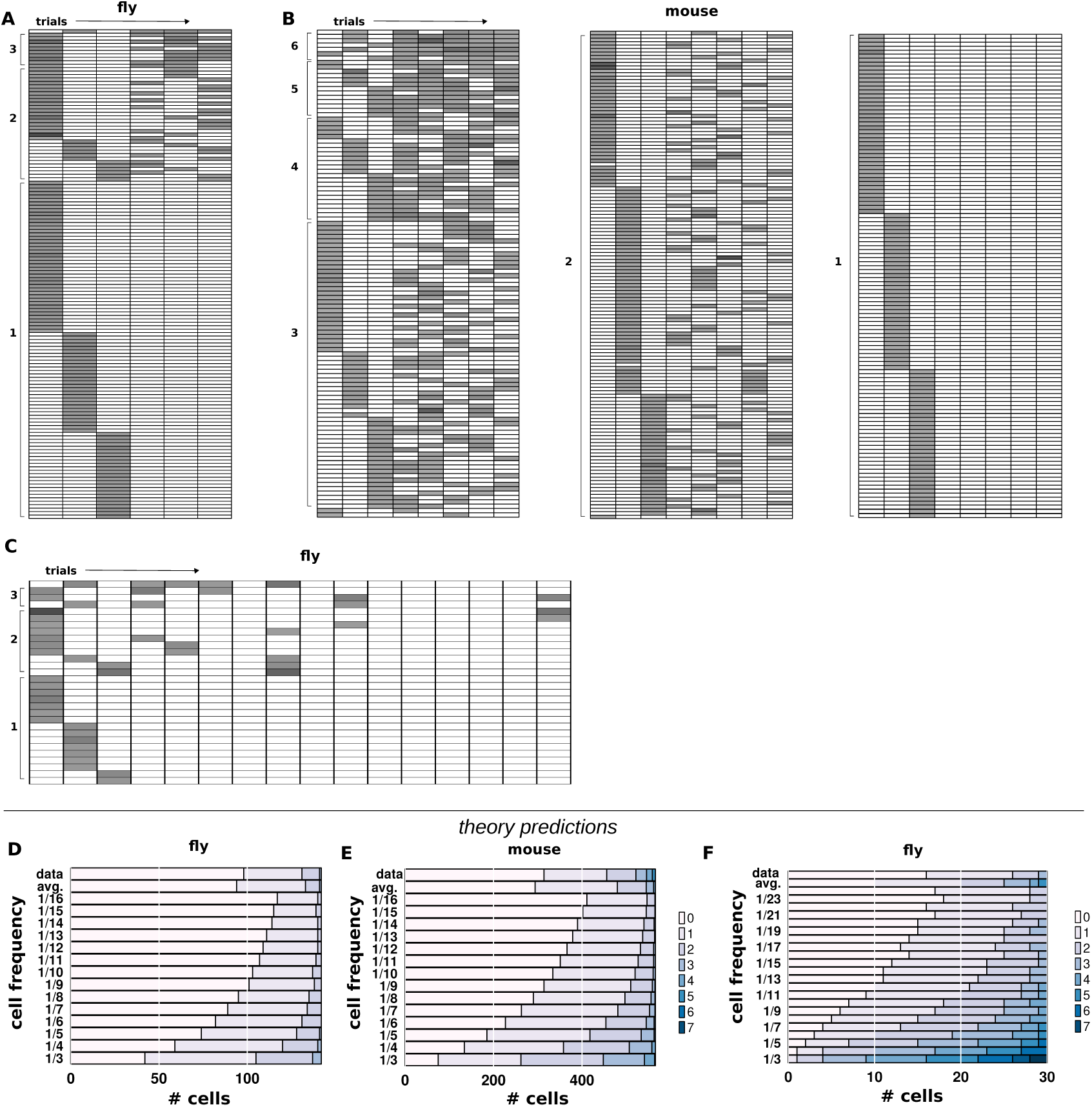
Unreliable cells are a composition of cells with different levels of reliabilities. (A-C) show stacked plots of responses to odors over 6,8, and 16 trials in the fly and the mouse for cells with only one response in the first 3 trials. Each row shows responses for a cell. In all three cases, cell responses in subsequent trials range over a spectrum of frequencies reflecting that cell reliabilities likely lie along a continuum. (A) Odor responses of 133 cells to 6 odors across 6 trials in the fly MB. The numbers on the left, in brackets, denote the number of cell responses over 6 trials. The rows are arranged in the decreasing order of number of responses. The rows are also sorted so that cells, e.g., with three responses, with a response in the first trial come first,followed by cells with a response in the second trial, followed by a response in the third trial. (B) and (C) are arranged similarly. (B) Odor responses in 542 cells in the mouse PCx across 8 trials. The last column showing a single response is shortened in the interest of space and represents 245 cells. (C) Odor responses over 16 trials in the fly MB in a total of 30 cells. (D-F) theoretical predictions for the number of cells of each reliability (from trial 4 onwards) for different cell frequencies. (D) Predictions of composition of cell reliabilities in the population for the fly data set with 6 odors and 6 trials per odor for a total of 142 cells. Going from left to right on the x-axis, the shades of blue from light blue to dark blue designate the number of cells that respond with that reliability (legend on right shows the color-frequency mapping). The bottom row shows the prediction if all cells have a base probability of response of 1/3, i.e., they respond once every 3 trials. For a cell population of 142 cells, we would have 42 cells that respond 0 times in trials 4 to 6, 63 cells that respond once, 32 cells that respond twice, and 5 that cells respond 3 times. The designations on the y-axis denote the different cell frequency levels, and the corresponding row, the cell composition. The top two rows show the average of the theory predictions and the data. For the fly, the top two rows are similar suggesting that cells in a population might be composed of different frequencies. (E) Predictions of cell numbers for different cell frequencies for the mouse data set with 10 odors, and 8 trials per odor, for a total of 544 cells. The bottom line/row represents the number of cells at each frequency for trials 4 to 8 if all cells had a base probability of response 1/3. For a cell population of 544 cells, we would have 75 cells that respond 0 times in trials 4 to 8, 187 cells that respond once, 186 cells that respond twice, and 93 cells respond 3 times, 23 cells that respond 4 times, and 2 cells that respond 5 times. The top two rows (like the fly) are similar reflecting that the cell population contains cells with a range base cell response probabilities. (F) Predictions for a fly data set with 1 odor, and 16 trials of the odor for a total of 30 cells. This plot is similar to (D) and (E) with the rows showing cell frequency compositions for cells with different base probabilities. Unlike (D) and (E), the average and data appear dissimilar, although the data does not resemble any particular cell base probability row, showing that it is likely that the cell population is made of a composite of cells with different base probabilities.

**Figure S7:**
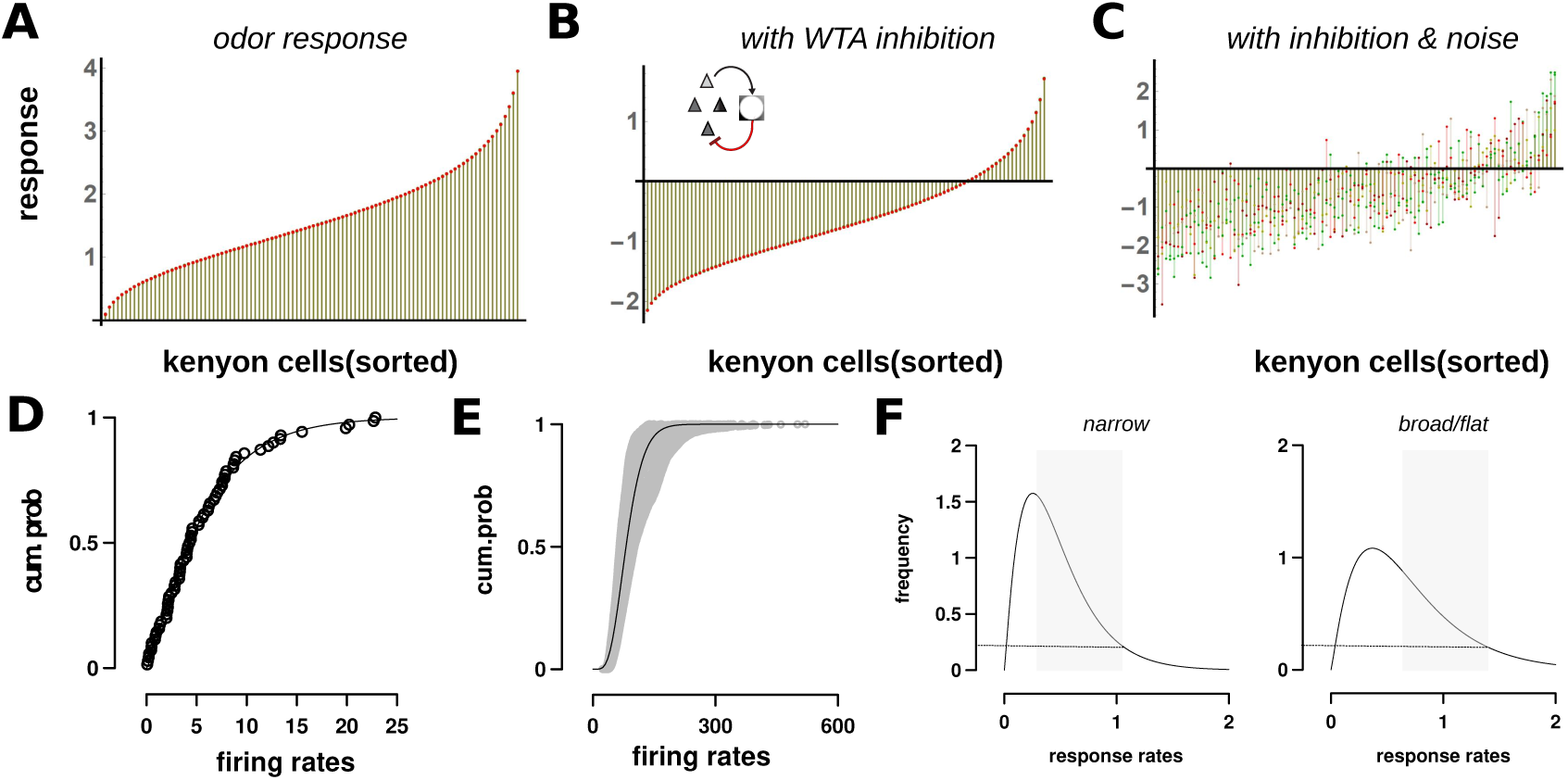
Noise in the winner-take-all mechanism produces a stochastic code. (A) A simulation of realistic odor response of KCs without APL inhibition, based on the data from Lin et al. (2014). The response distribution approximates a Gamma distribution and this plot shows KCs on the x-axis sorted in the increasing order of their response. (B) When subjected to APL inhibition fixed to be at 90th percentile of the Gamma distribution, all but the top 10% of cells are silent. The responses are the same from trial to trial as there is no noise. (C) When noise is injected into the feedback, we observe variable responses across all cells with three features. Cells in the top 5% are reliable and their average response is higher. Cells below 50% almost never respond, and cells close to the threshold are highly variable switching between responding and non-responding cells. (D,E) Cumulative frequency plots of odor responses in MB with APL turned off, taken from Lin et al. (2014), fit a Gamma distribution with shape=4.12 and scale=5.8. (D) shows an example plot for a single trial, while (E) shows the plot for about 70 different odor trials with the black line showing the (average) fitted Gamma distribution. (F) The breadth of the distribution influences the number of cells affected by noise. Compare the two Gamma distributions with narrow (left, shape = 0.22) and flatter distributions (right, shape = 0.32). The dotted line marks the top 10% of the cells. The width region marked in a shade of grey is equal to the noise amplitude, and shows that the region below the curve is much larger when the distribution is narrow. What this means is that more cells below the cutoff induced by the WTA circuit are influenced by noise, and thus there would be more unreliable cells.

**Figure S8:**
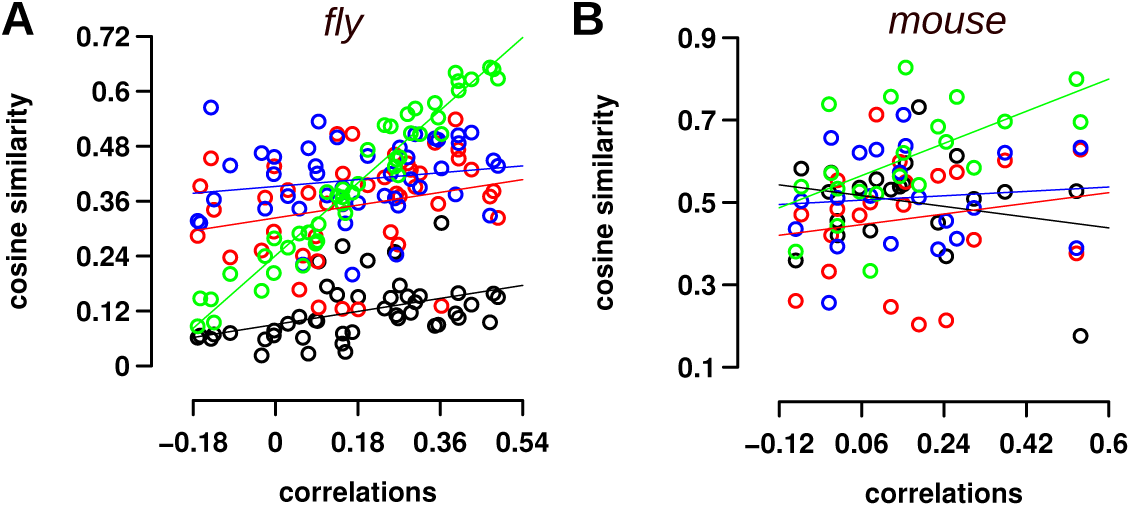
Sparse coding does not improve discrimination ability for similar odors. (A,B) Comparison of information about odor similarity for different populations of cells in flies and mice. (A) A non-parameterized plot showing how cosine similarity of four groups of cells changes with an increase in odor similiarity (as measured by Pearson’s correlation). In green are cells in the top 25% of responding cells, in blue are cells from 25 – 50%, in red are cells from 50 – 75%, and black designates cells in the bottom 25% of responding cells. The slope for cells in green (slope = 0.88) is highest showing that this population best captures the change in odor similarity, similar to reliable cells in Figure 4. It also shows that aggregate (over all odor pairs) similarity is highest for this set of cells. (B) The same plot for mice, showing that the top 25% of cells preserve more odor–similarity information, and that the similarity of odors does not change for the whole population compared to the top 25% of the population.

### Tables

**Table S1:** The top 40 parameter combinations (out of 44,217) of the noise exploration model that produced results most similar to the ones observed in Figure 1C,D. The first 4 columns are values of the parameters. Column 1: the APL gain that determines the WTA based sparse code, e.g., a value of 5 means that when there is no noise (deterministic system), the top 5% of cells are active. Column 2: PN noise describes the amount of noise at the PN level. Column 3: PN-KC: the amount of noise in the synapses between PN and KCs. Column 4: APL-KC: amount of noise in the feedback synapses from APL to KCs. In columns 2-4, noise is given as a fraction of the component’s value, and for percentage terms, it must be multiplied by 100. Thus, in the first row APL→KC noise is 0.275 of the synaptic strength or 27.5% of it. Column 5: The result of the optimization constraint, shows how far the model’s results are from observed results. The lower the value the better the fit. The table shows that all 40 parameter combinations contain noise in the APL→KC connections in the range 0.25-0.3. Correspondingly, PN and PN→KC noise range from 0 to 0.35, with quite a few entries being 0. Thus, noise in these two components is less important to producing the stochastic code than noise in APL→KC connections, which seem to be essential for producing observed results.

|  | APL gain | PNs | PN→KC | APL→KC | distance from observed |
| --- | --- | --- | --- | --- | --- |
| 1 | 7.00 | 0.00 | 0.05 | 0.275 | 0.11 |
| 2 | 8.00 | 0.05 | 0.00 | 0.30 | 0.11 |
| 3 | 8.00 | 0.10 | 0.05 | 0.30 | 0.12 |
| 4 | 7.00 | 0.20 | 0.20 | 0.25 | 0.12 |
| 5 | 7.00 | 0.15 | 0.00 | 0.275 | 0.13 |
| 6 | 8.00 | 0.15 | 0.10 | 0.30 | 0.13 |
| 7 | 7.00 | 0.15 | 0.05 | 0.275 | 0.13 |
| 8 | 8.00 | 0.00 | 0.35 | 0.275 | 0.14 |
| 9 | 7.00 | 0.35 | 0.10 | 0.25 | 0.14 |
| 10 | 6.00 | 0.15 | 0.05 | 0.275 | 0.14 |
| 11 | 6.00 | 0.20 | 0.05 | 0.275 | 0.14 |
| 12 | 7.00 | 0.05 | 0.00 | 0.275 | 0.14 |
| 13 | 8.00 | 0.05 | 0.15 | 0.30 | 0.15 |
| 14 | 8.00 | 0.30 | 0.05 | 0.275 | 0.15 |
| 15 | 6.00 | 0.05 | 0.10 | 0.275 | 0.15 |
| 16 | 6.00 | 0.25 | 0.00 | 0.275 | 0.15 |
| 17 | 8.00 | 0.30 | 0.15 | 0.275 | 0.15 |
| 18 | 8.00 | 0.25 | 0.00 | 0.30 | 0.15 |
| 19 | 7.00 | 0.00 | 0.00 | 0.275 | 0.15 |
| 20 | 6.00 | 0.15 | 0.10 | 0.275 | 0.15 |
| 21 | 7.00 | 0.30 | 0.15 | 0.25 | 0.16 |
| 22 | 7.00 | 0.00 | 0.10 | 0.275 | 0.16 |
| 23 | 8.00 | 0.30 | 0.00 | 0.275 | 0.16 |
| 24 | 7.00 | 0.40 | 0.05 | 0.25 | 0.16 |
| 25 | 8.00 | 0.05 | 0.05 | 0.30 | 0.16 |
| 26 | 8.00 | 0.00 | 0.10 | 0.30 | 0.16 |
| 27 | 7.00 | 0.05 | 0.10 | 0.275 | 0.17 |
| 28 | 8.00 | 0.35 | 0.20 | 0.275 | 0.17 |
| 29 | 7.00 | 0.15 | 0.35 | 0.23 | 0.17 |
| 30 | 7.00 | 0.30 | 0.00 | 0.25 | 0.17 |
| 31 | 8.00 | 0.35 | 0.00 | 0.275 | 0.17 |
| 32 | 7.00 | 0.10 | 0.25 | 0.25 | 0.17 |
| 33 | 6.00 | 0.00 | 0.10 | 0.275 | 0.17 |
| 34 | 6.00 | 0.05 | 0.05 | 0.275 | 0.17 |
| 35 | 8.00 | 0.20 | 0.25 | 0.275 | 0.17 |
| 36 | 8.00 | 0.00 | 0.30 | 0.275 | 0.17 |
| 37 | 7.00 | 0.15 | 0.10 | 0.275 | 0.18 |
| 38 | 6.00 | 0.10 | 0.05 | 0.275 | 0.18 |
| 39 | 7.00 | 0.20 | 0.25 | 0.25 | 0.18 |
| 40 | 6.00 | 0.00 | 0.35 | 0.25 | 0.18 |

**Table S2:** Results of the performance of 3 linear classifiers/decoders on the fly and mouse datasets: lda – linear discriminant analysis, knn – k nearest neighbor, svm – support vector machines. The decoders were trained for 3 types of cells – reliable (Rel.), unreliable (Unrel.), and all (all) cells – as described in Methods, and in this table we show the performance results in terms of accuracy (the fraction of the test set that was correctly predicted) and the averaged AUC measure. For all three decoders, it is easiest to distinguish between odor classes when only reliable cells are considered. Similarly, when all cells are considered, it is also more discerning than the unreliable class of cells, and except for the fly lda set, less slightly less discerning than reliable cells alone. Unreliable cells also have some amount of discriminatory power based on their non-negative accuracy scores, and high AUC scores. For all cases, the training set contained 80% of the odor trials, and the test set contained the other 20%.

|  | Accuracy |  |  | AUC |  |  |
| --- | --- | --- | --- | --- | --- | --- |
|  | Rel. | Unrel. | all | Rel. | Unrel. | all |
| lda (mouse) | 0.60 | 0.20 | 0.40 | 0.80 | 0.59 | 0.64 |
| lda (fly) | 0.57 | 0.28 | 0.50 | 0.72 | 0.72 | 0.83 |
| knn (mouse) | 0.40 | 0.50 | 0.10 | 0.74 | 0.39 | 0.59 |
| knn (fly) | 0.64 | 0.36 | 0.21 | 0.71 | 0.36 | 0.65 |
| svm (mouse) | 0.55 | 0.20 | 0.40 | 0.88 | 0.70 | 0.84 |
| svm (fly) | 0.64 | 0.14 | 0.42 | 0.93 | 0.66 | 0.82 |

**Table S3:**
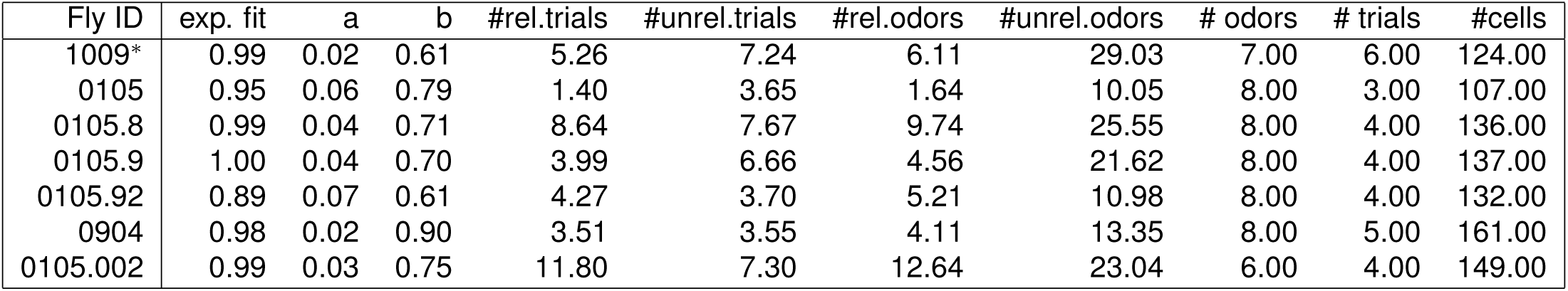
The response characteristics (Fig. 1) for all the flies that were examined. The first 3 columns give the fit of the exponential and the two parameters a and b of the fit equation, y = aebx. These columns are followed by the mean number of reliable and unreliable cells per trial and odor. The trials column gives the odor with the maximum number of trials. The double starred rows are ones where the total number of trials is lower than # trials * # odors. In order, they are 25, 22, and 35. As can be observed, the average trend is that the number of unreliable cells increases with total trials. The single starred fly (first row) is the one that was used for the figures in the main paper.

**Table S4:** The response characteristics (Fig. 1) for all the mice that were examined similar to Table S3. The first 3 columns give the fit of the exponential and the two parameters a and b of the fit equation, y = aebx. These columns are followed by the mean number of reliable and unreliable cells per trial and odor. Unlike the mouse, all mice experienced the same number of total trials, and as a result, they have similar numbers of average number of unreliable cells per odor. The starred mouse (first row) is the one that was used for the figures in the main paper.

| Mouse ID | exp. fit | a | b | #rel.trials | #unrel.trials | #rel.odors | #unrel.odors | # odors | # trials | #cells |
| --- | --- | --- | --- | --- | --- | --- | --- | --- | --- | --- |
| 164* | 0.99 | 1.10 | 0.49 | 3.40 | 8.66 | 4.42 | 43.09 | 10.00 | 8.00 | 285.00 |
| 163 | 0.98 | 1.09 | 0.42 | 1.30 | 7.04 | 1.67 | 38.65 | 10.00 | 8.00 | 318.00 |
| 7 | 0.98 | 1.72 | 0.36 | 1.62 | 9.66 | 2.31 | 49.12 | 10.00 | 8.00 | 147.00 |
| 8 | 0.96 | 2.34 | 0.42 | 2.44 | 6.60 | 3.14 | 35.54 | 10.00 | 8.00 | 121.00 |
| 9 | 0.98 | 1.77 | 0.40 | 2.39 | 10.30 | 3.16 | 51.10 | 10.00 | 8.00 | 209.00 |

## Methods

We split methods into three parts. The first part concerns experimental details, the second deals with analysis techniques used, and third describes the theory and modeling. For the theory and analysis parts, we used the statistical programming language R (Team, 2013), and the routines used for analysis will be made available at the time of publication, through Github.

### Experimental details

#### Mice

Experimental and surgical protocols were performed in accordance with the guide of Care and Use of Laboratory Animals (NIH) and were approved by the Institutional Animal Care and Use Committee (IACUC) at Brown University. C57BL/6J mice were crossed to Ai14 mice (Madisen et al., 2010) and male and female heterozygous transgenic offspring 8–12 weeks of age were used. Mice were maintained with unrestricted access to food and water under a 12–hour light/dark cycle and housed individually after surgery.

#### Stereotaxic Surgery

Virus (AAV1-syn-jGCaMP7f-WPRE, (Dana et al., 2019)) was purchased from Addgene and injected using manually controlled pressure injection with a micropipette pulled with a Sutter Micropipette Puller. Mice were anesthetized with Isofluorane with an induction at 3% and maintenance at 1-2% with an oxygen flow rate of *∼*1L/minute and head-fixed in a stereotactic frame (David Kopf, Tujunga, CA). Eyes were lubricated with an ophthalmic ointment and body temperature was stabilized using a heating pad attached to a temperature controller. Fur was shaved and the incision site sterilized with isopropyl alcohol and betadine solution prior to beginning surgical procedures. A 1.0mm round craniotomy was made using a dental drill centered to the following stereotaxic coordinates: ML: 3.9, AP: 0.3. Virus diluted to 1/3 in dPBS was injected at 100nL per minute at 3 different spots (total 1*µ*L) using the following coordinates (mm) to target piriform cortex (ML/AP/DV): 3.85/0.6/-3.8, 3.95/0.3/-3.9, 4.05/0.0/-4.0, all relative to bregma (Paxinos & Franklin, 2004). After 5 minutes, the micropipette was slowly retracted from the brain at 500*µ*m per minute. Following surgery, mice received buprenorphine slow release (0.05 – 0.1 mg/kg) subcutaneously. Lens implantation surgery occurred 2–3 weeks following virus injection. A GRIN lens (NEM–060–25–10–920–S–1.5p, GRINTech) was implanted above piriform cortex. The lens was implanted, centered to the craniotomy, at 100*µ*m per minute until reaching the following coordinate: DV: –3.9. Once placed, the lens was fixed to the skull with Metabond adhesive cement (Parkell, Edgewood, NY). A custom-made aluminium headbar was then attached to the skull using dental cement (Pi–ku–plast HP 36 Precision Pattern Resin, XPdent). Finally, a protective cap over the lens was applied with Kwik–Sil silicone elastomer (World Precision Instruments, Sarasota, Fl). Mice were allowed to recover from lens implant surgery for at least 4 weeks prior to imaging experiments.

#### Odor delivery

Animals were habituated to the experimenter and head-fixation setup for 30 mins a day for at least two days before the imaging experiment. On imaging days, odor stimuli were delivered through a custom built 16-channel olfactometer (Automate Scientific, Berkley, CA) equipped with a mass flow controller that maintained air flow at 1 liter per min. The olfactometer solenoids were triggered by a Teensy 3.6. A vacuum was applied inside the two-photon isolation box to evacuate residual odors. For all experiments, mice were habituated to the two-photon head fixed setup for 10 mins prior to imaging. An odor trial lasted 30 seconds (10 seconds of pre-stimulus baseline, 1 second of stimulation, 19 seconds of post-stimulus acquisition) with inter-trial intervals of 30 seconds. Odor stimuli were presented in pseudo-randomized fashion and 8 presentations of each odor were performed in a session. Two odor panel were run as follow (diluted in mineral oil vol/vol): Odor panel 1: benzyl isothiocyanate 10%, cinnamaldehyde 10%, (R)–(–)–Carvone 10%, Dihydrojasmone 10%, Ethylene brassylate 100%, Ethyl decanoate 50%, galaxolide 50%, Isoamyl phenylacetate 10% Odor panel 2: acetophenone 1%, amylamine 1%, butyl acetate 1%, ethyl hexanoate 1%, 2–Isobutyl–3–methoxypyrazine 1%, *β*-Ionone 1%, 2,3–Pentanedione 0.1%, Valeric acid 0.1%. A photoionization detector (miniPID 200B, Aurora Scientific, Canada) was used to confirm reliable odor delivery.

#### Two-photon microscopy

A typical imaging experiment lasted *∼* 1.5 hours per mouse. Two-photon imaging of the piriform cortex was performed using an Ultima Investigator DL laser scanning microscope (Bruker Nano, Middleton, WI) equipped with an 8Khz resonance galvanometer and high-speed optics set, dual GaAsP PMTs (Hamamatsu model H10770), and Z–Axis Piezo Drive for multiplane imaging. Approximately 90–150 mW of laser power (at 920nm, from Chameleon Discovery NX Ti:Sapphire laser source (Coherent Inc, Santa Clara, CA)) was used during imaging, with adjustments in power levels to accommodate varying signal clarity for each mouse. After focusing on the lens surface, optical viewing was switched to live view thru the two-photon laser, and a field of view (FOV) was located by moving the objective *∼*100-500 *µ*m upward. 3 FOVs were chosen separated approximately 80 *µ*m apart in depth. Images were acquired with a Nikon 10X Plan Apochromat Lambda objective (0.45 NA, 4.0 mm WD). GCaMP7f signal was filtered through an ET–GFP (FITC/CY2) filter set. Acquisition speed was 30Hz for 512 x 512 pixel images. Planes were imaged simultaneously, yielding a final acquisition rate of 4.53 frames per second.

#### Imaging data processing

Suite2p (Pachitariu et al., 2017) was used for video non-rigid motion correction, cell region-of-interest (ROI) selection and calcium trace extraction. Suite2p was run separately for each plane. Putative neurons were identified, and sorted by visible inspection for appropriate spatial configuration and Ca^2+^ dynamics. Neuropil signal was subtracted by a coefficient of 0.7.

## Data analysis

The methods used in analyzing the fly and mouse data are described below. To make it more accessible, we break it down by figure.

### Figures 1 & S1: analysis of response characteristics

For both flies and mice, all trials were interleaved, i.e., an example run of the first 8 trials might have looked something like this: odor1, odor5, odor6, odor3, odor4, odor8, odor3, odor1… For each trial, Ca^2+^ responses were recorded for the entire trial length that comprised three periods: the first period had no odor, a short odor exposure, and then a long period with no exposure. The first period for mice lasted 10s, and for flies also it was 10s. Following that, in both cases, odor exposure lasted 1s, and then a long period of no exposure. The odor response was recorded as the mean responses for the 2.5s following odor exposure (including the 1s odor exposure period).

To determine if a cell’s response was significant, we first calculated the mean and standard deviation of the odor response values for the first period, i.e, the background period without any odor exposure. A cell was considered to have had a significant response if its odor response value was 2.33 standard deviations (background) above the background mean. Figure 1C shows the analysis of odor responses of 124 KCs across 42 trials of 7 odors. The *y*-axis measure of the percentage of cells represents the number of significant cells divided by total number of cells. The adjoining graph gives the averaged numbers for all trials for each odor. Figure 1E, plots the reliability of each cell-odor pair versus the mean response size. The mean response size here is the expected value, i.e., Ʃ*_trials_* prob. of response(trial*_i_*) *∗* response.size*_i_*.

The calculations for the mouse PCx responses were similar. The mouse data in the main paper shows the responses of a set comprising 10 odors (2 controls) and 8 trials per odor, for 285 PCx cells. Here, too, odors were interleaved across trials. A response was considered significant if the mean response for the 10 time frames since odor onset was greater than 2.33 SDs above the background mean. Both SD and mean for the background were measured for the approximately 10s before odor onset. The length of a time frame is 0.453 s.

For Fig. S1A,B, we used a linearized form of the exponential equation, log(*y*) = log(*a*)+*bx* to get the quality of the fit, whilst making sure that log errors were propagated properly. The exponential forms of fit equations are shown in Figure 1E,G.

### Figure S2: noise varies with signal size, supplemental item for Figure 1

The presence of trial-to-trial variability led us to wonder if the two sub-populations of cells could be differentiated not only on the basis of how frequently they respond but also on the variability of their signal. We had shown earlier that signal and frequency of response are linearly correlated. We next wondered if this relation might also extend to the noise or variability of signal across trials. Studies have used two measures of noise, coefficient of variation (CV) and the Fano Factor (FF). While CV is defined as the SD over the mean, fano factor is variance (or SD^2^) over the mean. For a Poisson distribution, wherein the variance is the same as the mean, FF would be 1.

Figure S2 captures the relation of the signal to the noise through the CV and FF measures. Each grey dot represents the FF and CV for a single cell for one odor. The CV and FF measures for flies and mice were below 1. A significant difference between flies and mice was that in the fly, CV and FF measures were indistinguishable by cell frequency or a cell’s reliability. In the mouse, on the other hand, cells with a lower trial frequency or reliability had a higher CV or FF measure. This finding reflects the fact that while in the fly, increases in average signal size kept track with a cell’s SD (Fig. S2), in the mouse, although the two quantities were positively correlated, the rate of increase in SD in relation to signal size was less (Fig. S2). Thus, in flies, for most levels of reliability, there was no significant difference. In mice, however, cells with the highest reliability, had a lower level of CV and FF indicating that they might carry more robust information than other cells.

### Figures 2 & S5: analysis of response, overlap, and reliability distributions

We will use the the cumulative frequency distribution of reliabilities as an illustration of how we made the cumulative plots. We first sorted all reliabilities in increasing order. We then assigned a value of one to the lowest reliability, and for each subsequent item we added 1. If a bunch of cells had the same reliability, say 3, we added the number of these cells to the counter. The counter keeps track of the number of cells at or below this reliability value. Finally, we divided this counter by the total number of cells to get the cumulative frequency histogram values, and plotted them on a graph as shown in Fig. 2B. The procedure we outlined is a non-parametric way to plotting frequency histograms, explained elegantly in (Newman, 2005)

### Figures 4, S3 & S4: analysis of correlations and overlap between odors

For the first part, to compare the similarity of odors in antenna and MB, we used Pearson’s correlation coefficient. For the antenna, we calculated the correlation between all the odor pairs that were common to that dataset (Hallem et al., 2006) (for OSNs) and (Campbell et al., 2013) (for KCs). We then computed the correlations for the KC responses of these odor pairs. For the KC correlations of each odor-pair, we computed the correlation between 36 trial pairs (6 trials for each odor), discounted the same trial odor-pairs, and took the mean of the 30 odor-pairs to be the correlation. This KC correlation was on the y-axis. Note, that the odor-pairs shown in Fig. 4 are restricted to those whose correlations in the antenna and MB are within 0.2 of each other, since the Antenna responses are single trial responses and thus, might be more noisy than data collected over 6 trials in MB. Figure S3 shows that the qualitative results are the same even when all odor-pairs are considered. We also had access to 2 more datasets (with 5 more odors besides the main dataset), and we calculated correlations similarly for those datasets and have included them in this analysis (Figure 4A-B).

Figure 4E-H gives the overlap between similar and dissimilar odor-pairs. For this analysis, we only used the main dataset. First, how did we determine similar and dissimilar odors? Figure S5 shows the distribution of odor similarities using correlation as a similarity measure. The correlation shown here is the correlation between the average response vectors, i.e., responses averaged across all trials. Odor-pairs that had a correlation less than 0.18 were considered. In essence it was 0.15 as there was no difference between 0.15 and 0.18. We chose 0.18 because when you choose vectors from a normal distribution and compare their correlations, 95% of them have a correlation *≤* 0.18. Similarly, for the lower limit of similar odor pairs, we chose 0.35, as to isolate a cluster of odors.

For every odor, we divided cells into their reliability classes, and calculated the cell’s probability of response, which is the number of significant responses/total number of trials. Now, for the two odors A and B in an odor-pair, we can calculate the probability of the cell responding to A or B, and overlap is the probability that it responds to both, which is p(A).p(B). When we are considering odor-pair AB, the cell’s overlap would be the reliability of the cell for odor A. The plots show the overlap calculated this way for every cell in each of the odor-pairs.

Figure S3 gives the overlap vs reliability measures for cells across all odor pairs, not just similar or dissimilar ones. In this figure, we can see that the average overlap over all odors, reduces the overlap of similar odors. This is because the overlap for a cell, when odors are dissimilar is low across all reliability classes (Fig. 4G,H), and thus reduces the average overlap for cells with high reliability compared to the case of similar odors alone.

### Figures 5 & S8A, B: analysis of sparse coding

We compare 4 progressive groups of cells: the top 100, 75, 50, and 25 percentiles, to examine if these groups are more alike between similar odors. Here, when we say top percentile, we mean top percentile of responding cells. In flies, this would be roughly around 9 cells (since top 25% of 30 is around 8), and in mice, it would be around 28 cells.

Consider 2 similar odors, and that we would like to compare how similar the top 25% of cells for these two odors are. As a first step, we rank order the expected value of the odor response of each cell. We take the top 25% of cells for each odor, which could range from around 8 to 12 cells, because there will be some cells that respond to the both odors, while others do not. We then compute the cosine similarity of the average value of these cells for the 2 odors, and this value represents one grey point in the plot. Similarly, we compute the cosine similarity for all similar and dissimilar odor-pairs for each cohort of cells shown in Figures 5A,C. Here, we restricted dissimilar odors to include only those odors whose correlation (of average values) was less than 0.15. Similar odors had a correlation greater than 0.5. The number of dissimilar odor-pairs are fewer here compared to Figure 4, because in it we used an average of trial correlations, while here since we used average values for computing cosine similarities and percentiles, we also used average response values to compute correlations. The correlations of average response values are greater than the average of correlations across trials.

For Figures 5B,D, we took the top 25% and bottom 25% of responding cells for an odor, then we assigned them to the appropriate cell classes and counted that number. The points in grey are the numbers for the top 25% of cells, and the points in red are the numbers for the bottom 25% of cells for each odor. The averages are shown in black and darker red. And, we followed the same procedure in anaylzing the fly and mouse cells, except for the mouse we only considered the top and bottom 10% of cells: since the absolute number of mouse PCs cells at 10%, was equal to the number of KCs at 25%.

For Figures S8A,B, we computed the cosine similarity amongst the four quartiles of size 25 – top 100-75, 75-50, 50-25, and 25 – for all odors. For both, flies and mice, the 25% quartile captures odor similarity information faithfully. When odor similarity is low, cosine similarity is low, and increases with increase in similarity of the population, indicating that they faithfully carry odor similarity information. The other three quartiles have flatter slopes indicating that they carry less information that can distinguish odors.

### Figures 4 & S3D: analysis of reliable, unreliable, and all cell responses with machine learning algorithms

In order to assess the ability of unreliable and reliable cells to classify odors, we isolated cells of each type, and then fed them to a decoder (based on simple machine learning algorithms) to examine their ability at classification. If either of the cell types were to carry odor distinguishing information, then the decoder should be able to correctly classify odors when using these cells. We examined odor responses to three classes of cells: reliable, unreliable, and all significantly responding cells. To illustrate how we used the cells, we will use reliable cells as an example.

For each of the trials from an odor, we isolated all those cells that were reliable for that odor, and set the values of all other cells to 0. We repeated this procedure for every odor. Keep in mind that a reliable cell for one odor, might be unreliable for another, and thus would be positive for the first odor while being set to 0 for the second odor. There were some cells that were either silent or unreliable for all odors, and did not play a part in classification. We then fed these updated odor responses to a simple decoder. For decoder classification, we split the data set into 2 groups: a training set and a test set. For all groups (reliable, unreliable, and all cells), the training set contained 80% of trials, and the test set contained 20% of trials. For most of the decoders, we used a training set with trials chosen equally from all odors. The remaining 20% was used for testing. Finally, to ascertain that the results did not arise because of random effects on how the test and training sets were chosen, we used *k*-fold cross validation (Hastie et al., 2009), in which we set fold size to 20%. At the end, the presented results were averaged.

Classifier performance was reported using two measures: accuracy, and multiple class area under the receiver operator curve (AUC) according to Hand & Till (2001). Both measures were calculated on the test set. Accuracy is the fraction of trials in which the predicted odor and the actual odor are the same. The second measure, which we call all-to-all (A2A) AUC, is computed for all pairs of odors, and the numbers plotted in Figure 4 are the average AUC for all odor pairs. The individual pairwise odor AUC was calculated as the average of *A*(*i|j*) and *A*(*j|i*), where *A*(*i|j*) is the probability that a random trial of odor *j* has a lower probability of being classified as odor *i* than a random trial of odor *i*. A high score (maximum 1) denotes that the two odors *i* and *j* are easily separable. A low score indicates a high likelihood of them being mistaken for each other. We used 3 types of machine learning algorithms (2 with a linear kernel and one non-parametric): linear discriminant analysis, K-nearest neighbors, and SVM or support vector machines with a linear kernel (Hastie et al., 2009). The scores and the training:test set ratios are given for each of the decoders in Table S2.

As a test case for the A2A AUC, we also computed AUC values for odor-pairs that were either similar or dissimilar (Fig. S3D). We found that similar odors had a low average AUC while dissimilar odors had a high average AUC. The AUCs depicted in Fig. S3D were calculated for all cells. For this calculation, the classifier that we employed was kNN on the mouse dataset collected in this paper. We observed similar results with the linear SVM classifier and the fly dataset.

## Modeling and Theory

### Figure 3: WTA mechanism responsible for a stochastic code

We used a linear rate firing model of the olfactory circuit from the antennal lobe to MB, similar to previous models of the olfactory circuit (Srinivasan et al., 2018; Srinivasan & Stevens, 2019). In the antennal lobe, we mimic odor activity of PNs by sampling from an exponential distribution, based on analysis (Sterling & Laughlin, 2015) of findings from previous studies (Bhandawat et al., 2007). For the connectivity matrix from PNs to KCs, we used the same parameters revealed by an analysis (Sterling & Laughlin, 2015) of data from Caron et al. (2013). We generated the connection matrix in the following way: For each KC, we generated the number of claws it has by sampling from a Poisson distribution with *λ* = 0.85, and *n* = 8. Then, each of these claws would get connections from one of 50 PN types, and the PN was chosen by sampling from a hypergeometric distribution with parameters *n* = 50, and *a* = 0.07. Finally, for the strength of the synapses, we sampled from a Gamma distribution with shape and scale factors of 4 and 4. For the next step of the WTA involving KCs and APL, we used data from Takemura et al. (2017) and Li et al. (2020), who showed that in both directions, KCs *↔* APL, the number of contacts was mostly around 10 or greater. In this case, the connection strength in both directions is a convolution of Gamma distributions. This would be a Gamma distribution again, except in this case, with a much smaller standard deviation compared to the mean. In such a case, these two parameters (of connectivity strength) have less significance for responses levels of KCs as the gain of the APL*↔*KC loop is the important parameter, not the synaptic strength. For this reason, the WTA circuit parameters were chosen so that around 8% of KCs had a response, and later we also explored the effects of changing the gain of the loop.

*Noise*: To examine the effects of noise on the circuit, we introduced noise within various components, similar to previous investigations of the olfactory circuit (Srinivasan et al., 2018; Srinivasan & Stevens, 2019). For each component, we added multiplicative noise sampled from a normal distribution, e.g., if a PNs firing rate was *P_resp_*, the noisy version of the PN, *P_noise_* was *P_noise_* = *P_resp_*(1 + *η*(0*, σ*)), where *σ* designates the amount of noise. If the noise level was 10%, *σ* = 0.1. We explored noise in the following 4 basic (only in 1 component) scenarios: (1) Noise in the PNs (Panel C), (2) Noise in PN*→*KCs (D), (3) Noise in KC *→* APL connections (E), (4) Noise in APL *→* KC connections (F).

The parameters used for the different models in Figure 3C-G are as follows. For all the models, the number of PN types was 50, and the number of KCs was 150. The number of odors was 6, with 6 trials per odor. The APL inhibitory feedback was set so that only the top 10% of cells were active without noise present, and only the top 8% were active for model G. These two parameters were chosen as produced the desired reliable/unreliable cell ratio for the least amount of noise. The data plotted in the graphs is the mean of 5 runs. The error bars represents standard deviations.

The individual parameters for noise for each model. (C-F) Range of 0 to 1, in increments of 0.1. (G) Range of 0 to 0.5, in increments of 0.025.

We used the following parameters for the PN*→*KC connectivity matrix. For each KC, to determine the number of its claws, we sampled from a Binomial distribution with n=8, and p=0.715. Each claw connected to PN numbered from 1 to 50 based on a discrete hypergeometric distribution with a paramater of 0.07, and the connection strength was given by a Gamma distribution of shape and scale factors of 4 each.

### Figure S7: additional exploration of WTA circuit

In panels A-C, we mimicked realistic responses to odors in the absence of APL inhibition, i.e., the basal response without WTA, by first fitting data from Lin et al. (2014) that was generously shared with us by Andrew Lin and Gero Meisenbock. We found that the response fit a Gamma distribution with shape = 4.12 & scale = 5.12 (shown in panel E), and panel A shows the response of a 100 cells drawn from this distribution and sorted by increasing order of response size. The next panel shows the response sizes, wherein the WTA inhibition is set to the top 10*^th^* percentile. Panel C shows the same responses, except this time the inhibitory feedback has multiplicative noise added to it with an amplitude of 20%.

In Figure 2, we showed the response sizes of of all odor-KCs pairs from every trial. Here, in Panel D, we show the responses for one sample trial, again fit to a Gamma distribution with shape=1.12, and scale=0.32.

Panel F shows the probability distribution function for 2 curves with Gamma Distributions, with shape = 0.22 (left) and 0.32 (right). The change in shape makes the distribution narrower from the left panel to the right panel.

### Table S1: exploration of noise in a model of the fly olfactory circuit

Table S1 lists the fly model parameters that give the closest results to observed data. It shows that the WTA mechanism is essential to the stochastic code that we observed in Figure 1. In order to determine the optimal parameters, we took four of the parameters of the fly circuit: the APL gain that determines the number of KCs that are active, noise in the firing rates of the PNs, noise in the connection synapses between PNs and KCs, and noise in the inhibitory feedback from APL to KCs. We did not include noise in the connections from KCs to APL, because as we showed in Figure 3, even noise at the level of 400% does not produce observed results. This KC*↔* APL connection has little effect on the stochastic code. We did not vary other parameters such glomerular rates or glomerular*→*KC connections as we used empirical values that were uncovered in Bhandawat et al. (2007); Caron et al. (2013) and described in (Stevens, 2015).

To assess how well the model – with any particular combination of the 4 parameters – fits observed data, we exhaustively explored parameter space, while constraining the model with the 5 conditions observed with data 1: ratio of reliable/unreliable cells, number of reliable cells/trial, number of unreliable cells/trial, number of reliable cells across all trials, number of unreliable cells across all trials. Figure 3 provided us with two characteristics of each individual parameter. First, each parameter had a monotonic effect on these constraints, i.e., increasing the parameter either increased or decreased these constraints. Second, it gave us the maximum value that yielded observed results. These two characteristics led us to using the following parameter range in our explorations: APL gain from 5 to 13 in steps of 1, noise in PNs from 0 to 80% of firing rates in steps of 5%, noise in glomerular*→*KC connections from 0 to 80% in steps of 5%, and noise in APL*→*KC connections from 0 to 40% in steps of 2.5%. Thus, in all, we explored 44,217 parameter combinations. And, we ran each parameter combination 10 times, and took the mean of the results.

For each parameter exploration, we judged its fit to observed data in 2 steps, similar to an *l*1 *− norm* minimization procedure (Boyd et al., 2004). First, the model had to produce results for each constraint that fell within mean *±* sem range of observed results. Parameter combinations that did not satisfy this condition were discarded for the next step. Second, for each of the constraints we took the absolute value of the difference between the model’s and observed results normalized to observed results. For instance, for the constraint of reliable/unreliable cells, the ratio observed in experiments was 0.72, and what we observed with the parameter combination that was the global minimum (APL gain = 7, glom. noise = 0, glom-kc noise = 5% or 0.05, APL-KC noise = 27.5% or 0.275) was 0.712. So, the result for this constraint was (0.72-0.712)/0.72 = 0.01. Normalization was necessary in order to ensure a comparison between different constraints. For instance, if we consider another parameter, like the number of unreliable cells across all trials, its value for observed data is around 30. Thus, it is an order of magnitude higher than 0.72, the reliable/unreliable ratio constraint, and normalization is necessary for comparison.

The parameter combinations listed in Table S1 are the top 40 combinations arranged in increasing order of their distance – shown in column 5 – from observed results. The first combination is the global minimum. In all, out of 44,217 parameter combinations, 1290 yielded viable results, i.e., they satisfied each of the constraints. Amongst these, of the top 200 parameter combinations, 173 of them had APL-KC noise in the range of 20 – 30%. By contrast the range for glomerular and glom-kc noise went from 0 to 80%, suggesting that APL-KC noise was more significant and essential for producing a stochastic code. Additionally, in cases where there was no APl-KC noise, the amount of noise in the other two parameters jointly was in the range of 75 – 110%, showing that abnormally high amounts of noise is required in order to generate the stochastic code.

### Figure 6: a model of learning and discrimination based on the fly olfactory learning network

The objective of this model was to examine the contributions of reliable and unreliable cells towards discrimination. To have a realistic estimate of how they would work in an actual system, our model closely mimicked the actual fly association network detailed in the main text accompanying Figure 6, and depicted in Figure 7. For more in-depth detail, please refer to Modi et al. (2020).

There are two elements to the model. The first is the contribution per cell, and the second is the learning in the network, which is done by plasticity at the synapses between KCs and MBONs. We approximate the fly MBONs to two types: approach and avoid; the real fly has about 15 MBON types falling into either the approach or avoid categories. Learning is implemented in two stages that reflect the amount of training considered: normal, which in the fly case would be around 12 trials, and extended, which would be about 100 trials. With normal training, we consider the discriminatory power of the cell using equation 1. Initially, we assign equal weights to the connections from every KC to both MBONs. These weights undergo a change depending on whether they are from reliable or unreliable cells. The change in weights is governed by equation 3. Since reliable cells respond consistently and with a high response value their weight approaches 0 very quickly, i.e., by the end of the normal training period. Unreliable cell synapses, for the same reason, undergo little change in their synaptic weights. The overall discrimination as shown in equation **??** is the sum of all cells, which would be D(*A, B*) = D(*A, B*)*_R_* +D(*A, B*)*_UR_*. Since reliable cell synapses (from KCs to either the approach or avoid MBON depending on whether it is punishment or reward) are close to 0, D(*A, B*)*_R_* is high, and as unreliable cell synapses are unaffected, D(*A, B*)*_UR_* is low. The contribution of D(*A, B*)*_UR_* is significant, however, since there are so many unreliable cells.

When learning is extended over 100 trials, there is no significant change in the synapses of reliable cells (they are already 0), but those of unreliable cells undergo a drastic change and are close to 0. Therefore the contribution of unreliable cells to discrimination is also high now. One might think that since equation 1 depends on the probability of response, it would be low. But, as a population, unreliable cell contributions are high as there are many more unreliable cells than reliable (29 compared to 5 for flies and 43 compared to 4 in mice). So, even if the unreliable cell responds in only 1 or 2 trials, there are at least 5 times as many cells, and their contribution is substantial.

For Figure 6, for easy calculation, we set the weights of all reliable cell synapses to 0 after normal training, and the weights of all unreliable cell synapses to 0 after extended training. Unreliable cell synapses remained nearly unaffected by normal training.

